# Deep Behavioral Phenotyping of Mouse Autism Models using Open-Field Behavior

**DOI:** 10.1101/2021.02.16.431500

**Authors:** Ugne Klibaite, Mikhail Kislin, Jessica L. Verpeut, Xiaoting Sun, Joshua W. Shaevitz, Samuel S.-H. Wang

**Affiliations:** Department of Organismic and Evolutionary Biology Harvard University; Princeton Neuroscience Institute princeton University; Department of Physics Lewis-sigler Institute for Integrative Genomics princeton University

**Keywords:** Autism, Mouse behavior, Open-field test, Computer Vision, Ethology, Markerless tracking, Behavior quantification, Posture, Dynamics

## Abstract

Autism is noted for both its genotypic and phenotypic diversity. Repetitive action, resistance to environmental change, and motor disruptions vary from individual to individual. In animal models, conventional behavioral phenotyping captures such fine-scale variations incompletely. Here we use advances in computer vision and deep learning to develop a framework for characterizing mouse behavior on multiple time scales using a single popular behavioral assay, the open field test. We observed male and female C57BL/6J mice to develop a dynamic baseline of adaptive behavior over multiple days. We then examined two rodent models of autism, a cerebellum-specific model, L7-Tsc1, and a whole-brain knockout model, Cntnap2. Both Cntnap2 knockout and L7-Tsc1 mutants showed forelimb lag during gait. L7-Tsc1 mutants showed complex defects in multi-day adaptation, lacking the tendency of wild-type mice to spend progressively more time in corners of the arena. In L7-Tsc1 mutant mice, failure-to-adapt took the form of maintained ambling, turning, and locomotion, and an overall decrease in grooming. Adaptation in Cntnap2 knockout mice more broadly resembled that of wild-type. L7-Tsc1 mutant and Cntnap2 knockout mouse models showed different patterns of behavioral state occupancy. Our automated pipeline for deep phenotyping successfully captures model-specific deviations in adaptation and movement as well as differences in the detailed structure of behavioral dynamics.

## 1 Introduction

Autism varies in its manifestations. Arising in early life, this neurodevelopmental disorder is defined by the behavioral and diagnostic criteria of abnormal social interactions and communication, stereotyped repetitive behaviors, and restricted interests [13,41]. Measured by this collection of traits, the intensity of individual components is highly variable. The entire range of variation is currently classified for treatment purposes as a single broad entity, autism spectrum disorder (ASD) [1,43]. In addition to cognitive and social variation, persons on the autism spectrum also express variable degrees of deficit in movement and sensory response [35, 36, 44]. Overall, the highly variable combination of phenotypes associated with ASD presents a challenge to classification and diagnosis.

For a highly heritable disorder such as autism, mice are an attractive model because they open the possibility of studying the consequences of a particular genetic or environmental factor repeatedly among many individuals. Mouse models of autism have been designed to reflect known risk factors and causes in humans (construct validity) [10,30]. These models are selected for investigation based on their signs that recall the human disorder (face validity), including perseveration, disrupted social preference, and deficits in flexible learning. Face-valid traits can be observed in conjunction with other traits that are considered of secondary interest, such as sensory and motor deficits. Those sensory and motor phenotypes originate from the same genetic perturbation that produced the primary traits of interest, making them potentially useful for linking symptoms to underlying mechanisms.

Traditionally, mouse behavioral testing consists of discrete measurements such as location in a multi-arm maze or a three-chamber test apparatus [12,14]. Such characterization omits both finer details of behavior and higher-order complex behavioral motifs [49]. These details can now be extracted efficiently using modern computational methods. Recent advances in automated tracking open the possibility of deep behavioral phenotyping consisting of simultaneous tracking of body-centric joint and body part positions, *x-y* position in an arena, and task performance, thereby providing a multilevel view of behavior [3,7,15,34]. This allows the measurement of both movement and cognitive/social features from a single dataset [20,21]. We developed a method using such a system for semi-supervised behavioral classification in individual mice during spontaneous activity. Our method describes long-term structure and fine-scale kinematic details in repeated recordings. Body part identification is consistent within individual mice over multiple sessions in the open field arena. From these detailed features we used posture-dynamical clustering [5,24,25,33] to quantify the entire repertoire of behavior.

We established a description of a commonly-used strain of mice, C57BL/6J (“Black6”), to characterize baseline behavior and to identify sex differences. Sex-dependent differences in gene regulation have been suggested to underlie the higher incidence of autism in males [17]. We then identified features of movement that were correlated with genetic background and specific genetic manipulations [14,27,40]. As a test of the method’s discriminatory power, we explored gene-knockout (KO) models of autism that perturb either the whole brain (Cntnap2 KO) or are cerebellum-specific (L7-Tsc1 mutant). These two lines are well-studied monogenic strains used to investigate autism endophenotypes [48]. Cntnap2 KO mice have been reported as hyperactive and have been shown to display a mild change in gait, along with reduced time spent with a novel partner in social behavior assays [8, 32]. Mice with Purkinje cell specific (L7) null mutation of tuberous sclerosis 1 (Tsc1) reportedly exhibit a variety of social deficits compared to wild-type littermates, along with changes in gait, increased time spent grooming, and decreased behavioral flexibility [9,45,46]. Postural defects have been recently observed in many mouse models of autism [40], revealing an opportunity for deep phenotyping using machine-vision methods. Both of these strains exhibit similar altered spontaneous and task behavior compared to wild-type when using coarse-grained metrics; we applied our deep behavioral phenotyping to test for distinct signatures of behavior under spontaneous non-task conditions.

## 2 Results

### 2.1 Deep phenotyping of open field recordings

We recorded over 700 trials in an open field arena (Fig. 1a) from 162 mice (Table 1). Each mouse was recorded for 20 minutes every day for a period of four days. We imaged the mice from below a transparent floor to allow observation of the paws and base of the tail (Fig. 1b). Movies were analyzed with a semi-supervised behavioral classification and behavioral labeling scheme (Figs. 1 and S1). In the first step, we used a deep neural network to measure the posture of each animal over time. Recent advances in body part tracking allow for the use of a small set of user-generated labels to train a neural network for labeling body parts [33]. We trained the LEAP network with 660 human-annotated images and achieved position estimates with median confidence probabilities ranging from 0.91 to 0.98 for the snout, chin, inner and outer limbs, and base and tip of the tail. Ears, body center and sides, and the tail center point were excluded due to lower estimation accuracy (median confidence probability of 0.81). This pose estimation step resulted in a time series of the two-dimensional position of each body landmark, *x_i_*(*t*).

**Figure 1:**
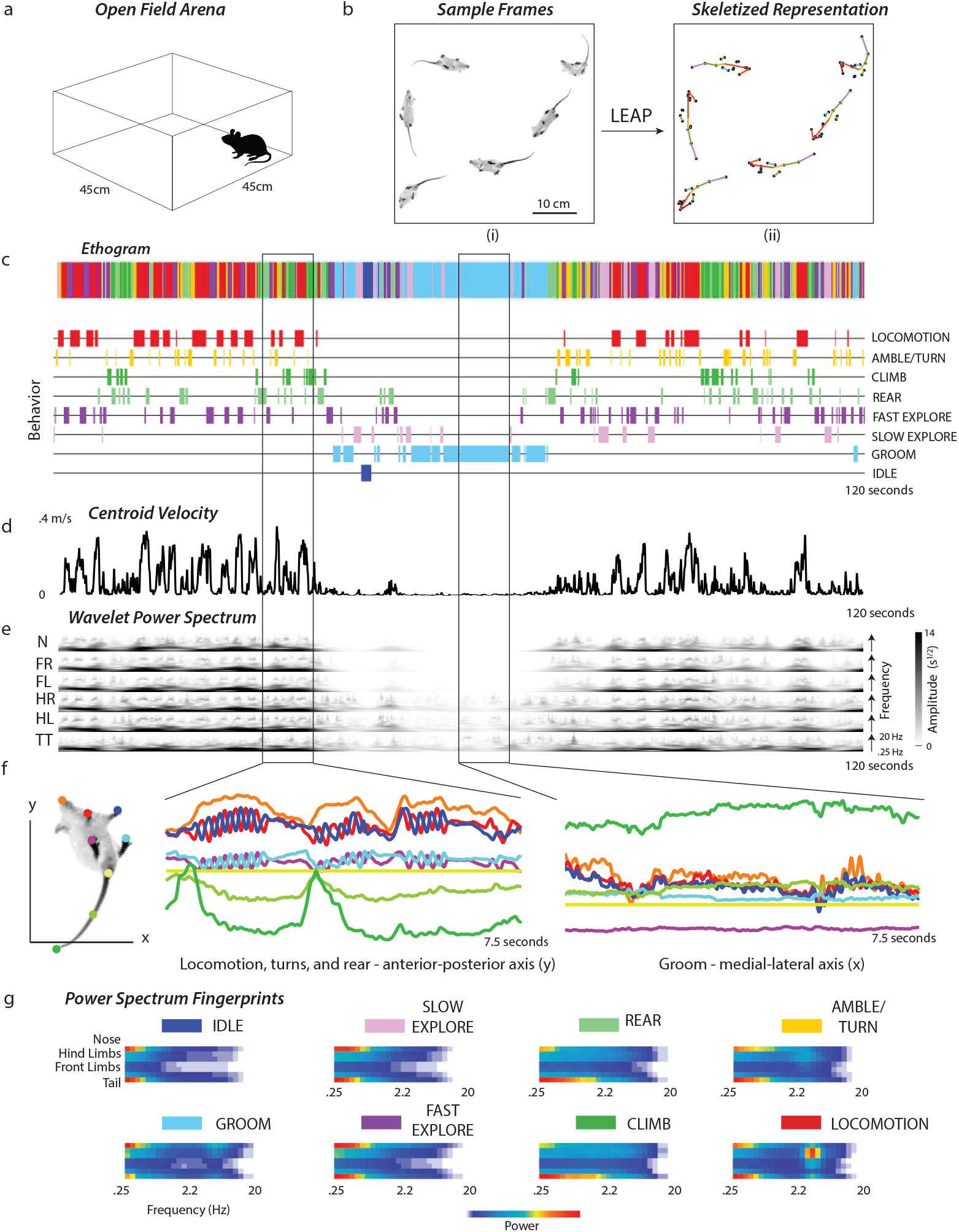
Processing of recordings in the open field produces multi-scale quantitative descriptions of behavior. a) Mice are imaged from below using IR illumination in an open field. b) A composite of several images from the open field recording. Both raw grey scale images (left) and skeletons obtained from virtual body markers labeled using the LEAP neural network (right) are shown. Scale bar is 10 cm. c) Ethogram of behavioral classes produced from our behavioral clustering and classification pipeline based on the posture and movement of animals over time. Below, the same time series is visualized as a raster to demonstrate behavioral usage during a two-minute period. d) Mouse centroid speed over the same two-minute period shown in panel (c). e) Raw power for select body parts. N: nose, FR: front right foot, FL: front left foot, HR: hind right foot, HL: hind left foot, TT: tail tip. f) Position time series for the nose, front and hind feet, and tail tip (left) during two behaviors. The *y* position, the anterior-posterior axis, is shown for a locomotion bout (center). The x position, the medial-lateral axis, is shown for a grooming bout (right). We use the full 2D position of each body part in our analyses, but show only the dominant axis for these behaviors here for brevity. g) Normalized power spectra for several tracked body parts for each of the eight behavior classes.

**Table 1:** Summary of experimental strains and number of mice recorded. Each mouse was recorded performing the open field test on four consecutive days.

| Strain | Number of Mice |
| --- | --- |
| C57BL/6J Males | 60 |
| C57BL/6J Females | 20 |
| Cntnap2 knockout Negative Littermates (male) | 14 |
| Cntnap2 knockout Heterozygote (male) | 15 |
| Cntnap2 knockout Homozygote (male) | 10 |
| L7-Tsc1 Negative Littermates (male) | 17 |
| L7-Tsc1 Heterozygote (male) | 17 |
| L7-Tsc1 Homozygote (male) | 9 |

The next step in our analysis was a semi-supervised clustering of postural dynamics. Our previous work on behavioral clustering analyzed the dynamics observed in whole images of an animal but did not specifically probe the dynamics of individual body parts [5,24,25]. We adapted this method for use with the body part position time series, *x_i_*(*t*) [33] and clustered the dynamics across timescales to define eight major classes of behavior in the open field (Figs. 1c, S1). The resulting ethogram revealed details not visible from centroid tracking alone (Figure 1d), namely the structure of spontaneous behavior across mesoscopic timescales.

The time series *x_i_*(*t*) was rotated and translated into the reference frame of the animal by aligning with the anterior-posterior axis defined by the snout and tail base body parts and transformed into the frequency domain using wavelet decomposition. This produced a high-dimensional output that captured multiscale (0.25-20 Hz) changes in posture over time (Figure 1e). We then performed *k*-means clustering of this high-dimensional data on a balanced selection of samples from recordings across all male mice. We initially clustered the sampled data into *k* = 100 clusters, enough to achieve finely-grained clustering that in some cases was not distinguishable by eye. Data from all experiments were then clustered by assigning each time point to the cluster of its nearest neighbors in the *k*-means training set.

An exploration of all 100 clusters revealed several broader classes of behavior exhibited by mice in the open field (Table 2). Similar to recent findings from fruit flies [4], we found that mouse behavior could be organized hierarchically with coarse classes composed of sub-classes representing variable aspects of behavior. For example, the 21 clusters that made up the ‘locomotion’ class represented the diversity of locomotion movements recorded, differing in velocity, amplitude and speed of limb movement, and the coordination of the limbs. We categorized each of the 100 clusters into one of eight behavioral classes (Figs. 1g and 2). This manual curation step revealed that time spent in the open field was composed of commonly studied behaviors such as locomotion and grooming, but also the less distinct movements mice made for a large fraction of their time including spatial exploration and ambling. We defined the classes *fast exploration* and *slow exploration* to characterize periods that are usually unaccounted for in traditional measurement paradigms. Fast exploration, for example, included quick turns and sniffs that are characteristic of alertness or anxiety, whereas slow exploration included head sweeps and extensions during calm periods.

**Table 2:** Eight coarse behavioral classes are used to describe the 100 fine-grained clusters obtained from a *k*-means clustering of the posture-dynamics signal. Descriptions were obtained by viewing movies sampling mice from each fine-grained behavioral cluster and grouping movies based on broad qualitative similarities.

| Class Label | Description |
| --- | --- |
| Idle | Mouse is still, no body parts are moving. |
| Groom | Variety of movements that involve repetitive motions, mainly using the front limbs and head, but also including tail grooming and hind leg scratches |
| Slow Explore | Slow head turning and sniffing |
| Fast Explore | Quick or directed movements of the head, sniffing, some forelimb placements |
| Rear | Moving into reared posture, slow hover on hind limbs |
| Climb | Reared posture with limb movement occurring mainly at walls and corners |
| Amble and Turn | Slow or uncoordinated locomotion that does not appear regular, single steps |
| Locomotion | Regular directed movement that involves limb coordination |

**Figure 2:**
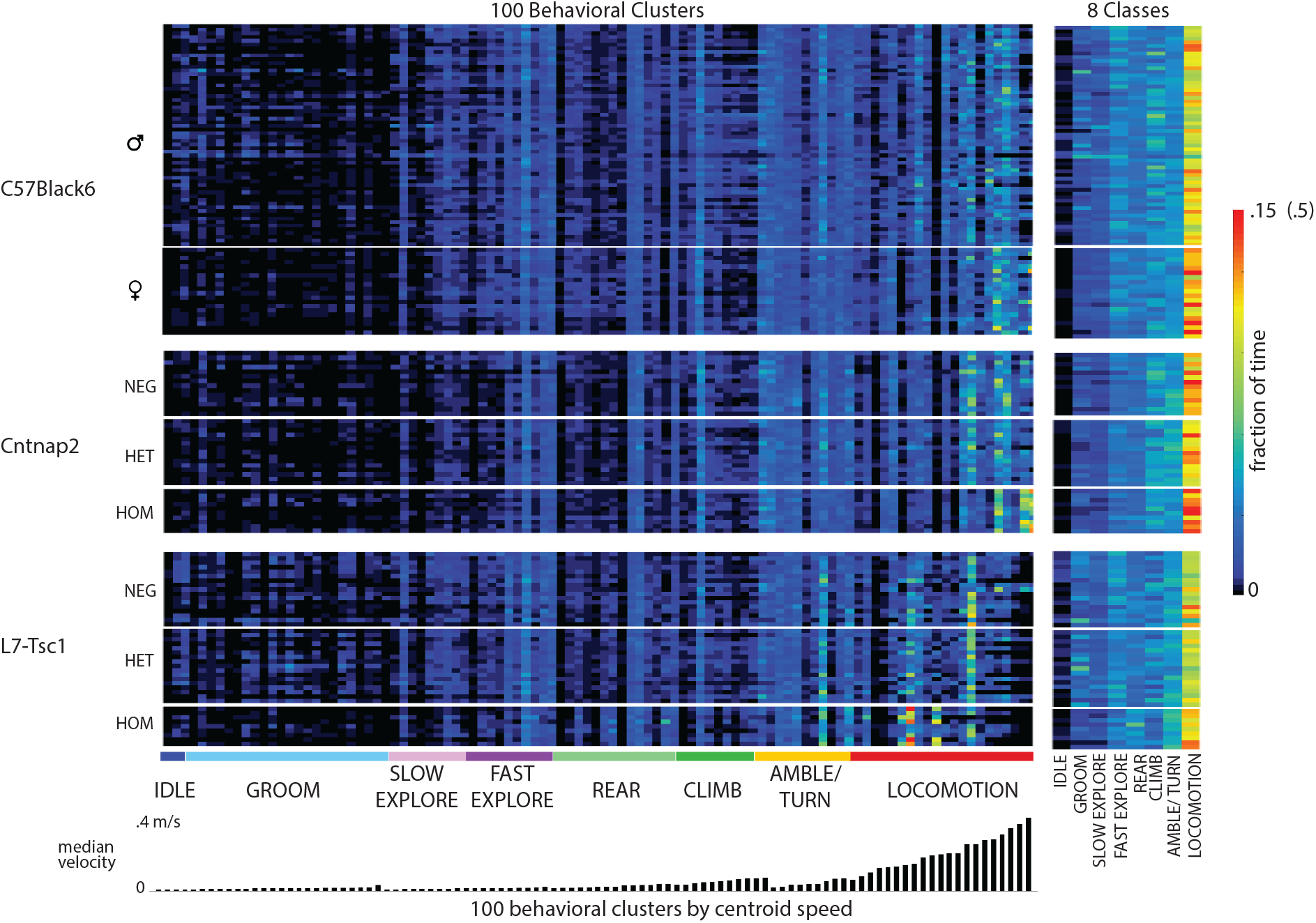
Behavioral cluster frequency across mouse models on Day 1. (left) Heatmap of the total behavioral occupancy of each behavioral cluster; rows are individual animals and columns are behavioral clusters. All rows within the 100-cluster matrix sum to one and describe behavior used over 20 minutes. Clusters are ordered by median centroid speed for each of the eight behavioral classes (bottom). (right) Heatmap of the behavioral occupancy for each of the eight behavioral classes.

We validated the accuracy and interpretability of these categorizations by visually inspecting randomly sampled movies of mouse behavior from each of the 100 clusters and the eight classes. Movie segments were sampled from all C57BL/6J male movies (300 20-minute movies overall), and limited to examples where the behavior in question was performed for at least 250 ms, or 20 frames at our frame rate of 80 Hz. Examples from the 100 behavioral clusters and eight behavioral classes are available in the Supplementary Materials.

### 2.2 Behavioral differences upon introduction to the open field arena

The fraction of time spent in each cluster in the experimental arena on the first day of exposure (Day 1) is shown in Figure 2. Clusters in Figure 2 are ordered horizontally according to their behavioral class, and then by median centroid speed within each class. Mice spent the most amount of time on Day 1 locomoting after placement in the novel arena, followed by turning, climbing, and rearing, but almost no time in the idle state.

Inspection of the 100-class usage across individuals revealed unique features that set the Cntnap2 KO and L7-Tsc1 mutants apart from their wild-type littermates, as well as from the large number of C57BL/6J mice, which had the same genetic background. C57BL/6J mice were most likely to use the fourth- and fifth-fastest locomotion modes at around .2 m/s on Day 1. The Cntnap2 KO was visually distinct from its corresponding wild-type and heterozygote littermates by the enriched usage of the several fastest locomotion clusters and a decrease in more moderate locomotion. This result was in agreement with findings that Cntnap2 KO mice are hyperactive [32] and accounted for the large distance these mice covered on the first day in the open field (Fig. S2). Full mutants from the L7-Tsc1 group showed the opposite trend from Cntnap2 KO mice, spending less time in the fast locomotion clusters than any other group. The L7-Tsc1 mutants often used several clusters of slow locomotion that were uncommon in either their control littermates or the baseline C57BL/6J mice.

### 2.3 Spatial habituation is reduced in L7-Tsc1 mutant mice

Mice, which are prey animals, tend to avoid open spaces in natural environments. Traditional analyses of open-field experiments divide the area into zones to quantify time spent in the open space, near the edges, and in the highly confining corners (Fig. 3). Mice spend most of their time at the edges and corners (thigmotaxis). Previous researchers have sometimes interpreted spatial preferences in terms of the emotional state of the mouse, often anxiety [2]. We found that in these corner positions, mice performed a variety of behaviors including grooming, rearing, climbing, and exploration (Fig. 3f).

**Figure 3:**
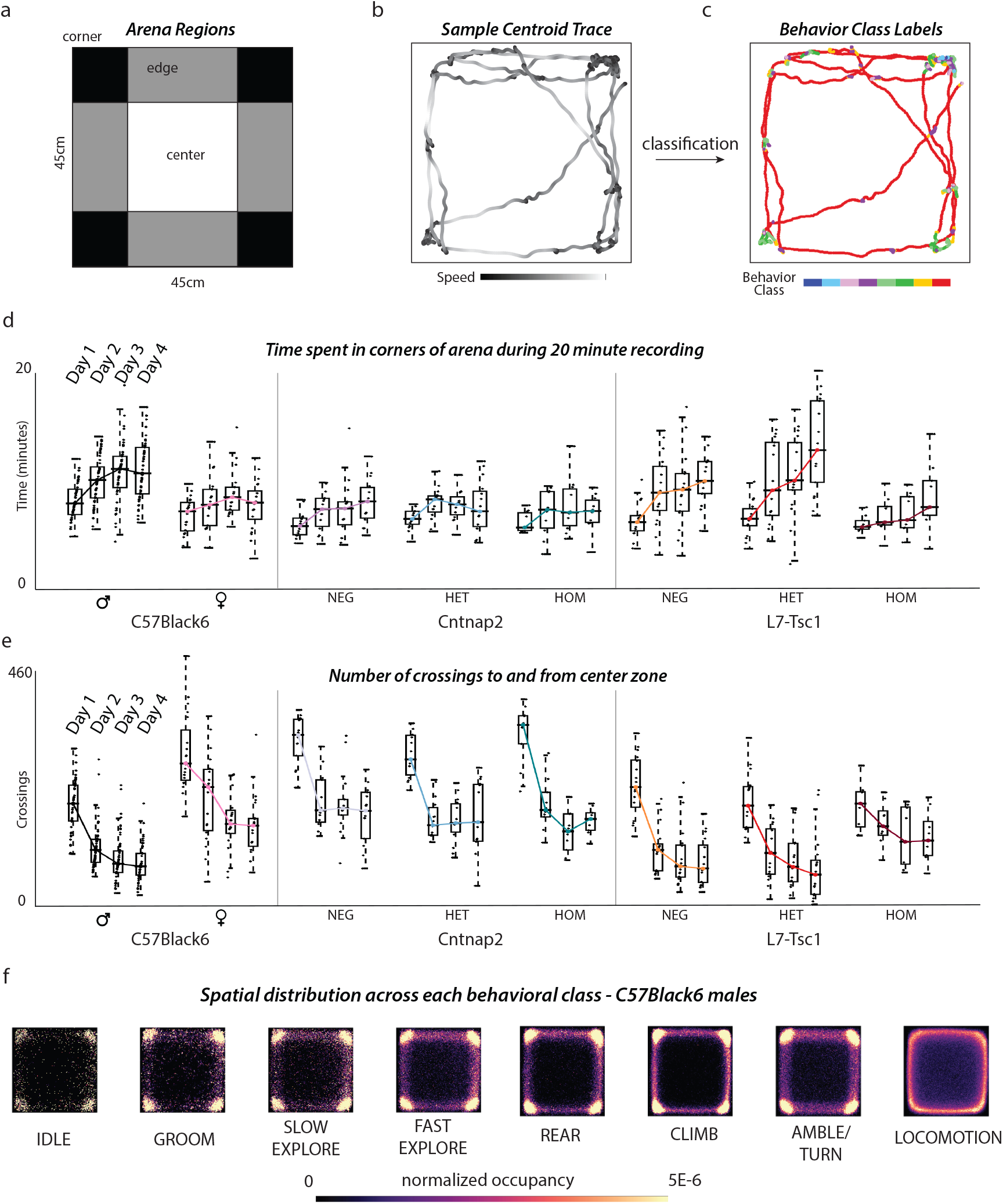
Mice perform behaviors in a spatially organized manner. a) Definition of the *center, edge*, and *corner* zones in the open field arena. b) The centroid trajectory for 120 seconds of activity in the open field. Grey scale represents the centroid speed for each time point. c) The same 120-second trajectory color coded by behavioral class. d) The total time spent in the corner regions for each mouse on each day displayed as a box plot. The median value is shown as a solid line in the color corresponding to the given condition. e) The number of center crossings for each mouse on each day displayed as a box plot. The median value is shown as a solid line in the color corresponding to the given condition. f) The spatial distribution is shown for each of the eight behavioral classes.

When first introduced to a new environment, mice explored the full space with some preference for corners. With time and repeated exposure to the same environment, mice habituated and spent less time crossing through the center of the arena and more time in the corners. We measured the time an animal spent in the corner zones of the arena for each of the 4 days of observation to identify the development of thigmotaxis. C57BL/6J males and females, as well as negative and heterozygote littermates of the ASD models, demonstrated a gradual increase over days in time spent in the corners (Fig. 3d). Cntnap2 KO mice did not show altered levels of spatial habituation within the statistical detection capacity of our experiments. In contrast, L7-Tsc1 mutant mice showed a significantly reduced level of habituation, with only a 26% increase in time spent in the corners between Days 1 and 4 compared to 34% and 46% for the litter mates. This reduced level of spatial habituation was also seen in the number of center crossings, where the mutants crossed 2.5 times more frequently than controls after four days of habituation (Fig. 3e). While L7-Tsc1 mutant mice showed a similar level of anxiety by these metrics compared to controls on Day 1 of the experiment, they were less capable of habituating to the new environment even after four successive days of exposure.

### 2.4 Behavioral habituation is reduced in L7-Tsc1 mutant mice

Mice also modulate specific behaviors as they habituate to a new environment. We found that both male and female C57BL/6J mice increased the time they spent idling, grooming, and exploring, and decreased the time they spent climbing, turning, and locomoting over consecutive days in the same arena. The largest shift occurred between the first and second day (Fig. 4a, S3a). Rearing stayed approximately constant from days 1 to 4.

**Figure 4:**
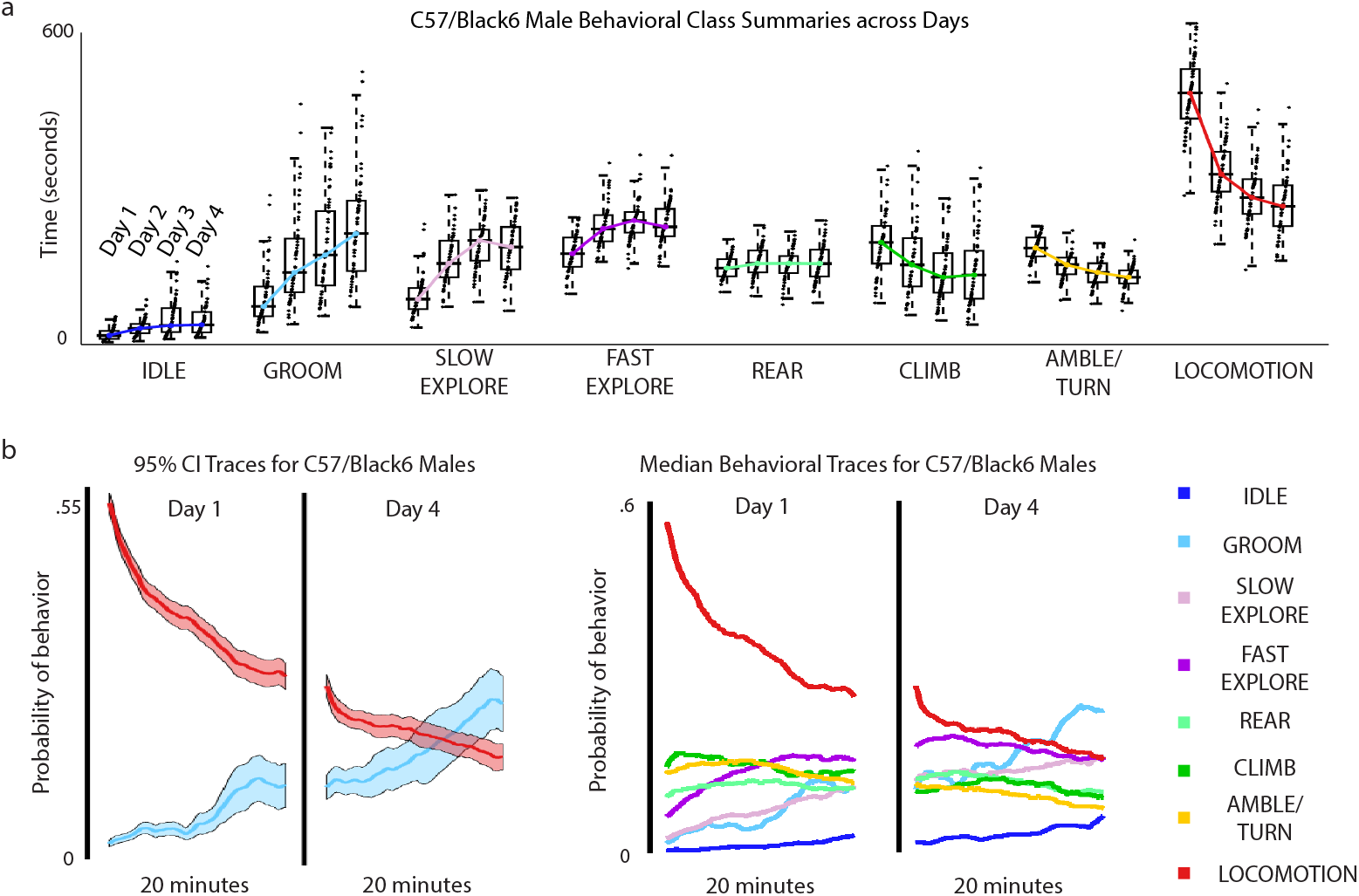
Behavioral usage over time for C57BL/6J mice. a) Behavioral usage for each of eight coarse categories plotted for C57BL/6J male mice for each of four observation days. All individuals are shown as points, colored traces correspond to the median fraction of time spent in the behavior for each day. b) The usage of grooming and locomotion behaviors during the 20-min observation period for Days 1 and 4. Shaded regions represent the 95% confidence interval. c) The mean usage for each of eight behavioral classes is shown for Days 1 and 4.

Behavior also changed within the 20-minute observation period of each experiment (Fig. 4b). The pattern of behavioral change exhibited by C57BL/6J mice within an observation period was similar to the across-day change: locomotion, climbing, and turning decreased over time while idling, exploring, and grooming increased (Fig. 4b). On average, males groomed for longer while females spent more time performing locomotion, with females in addition achieving higher speeds. A comparison of habituation curves for each of eight behavioral classes over four days of recording revealed this trend across all behaviors, with females performing active behaviors more at the onset of each experiment (Fig. S3).

Cntnap2 KO mice were unaltered in their ability to habituate and were not significantly different from their littermates although they did groom more often overall (Fig. 5). L7-Tsc1 mutant mice showed reduced habituation (Figures S3, S4, and S5).

**Figure 5:**
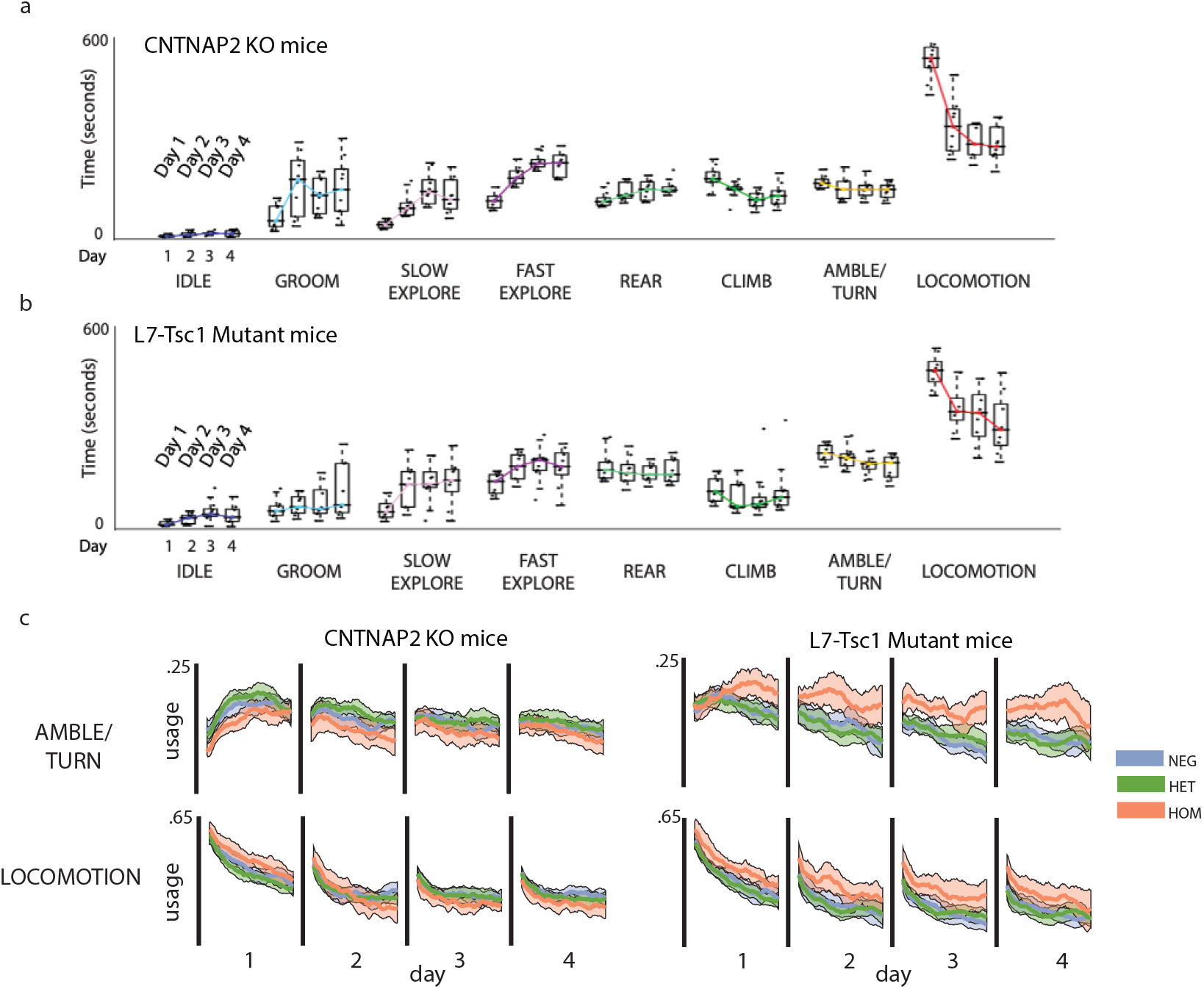
Behavioral usage over time for Cntnap2 KO and L7-Tsc1 mutant mice. a,b) Behavioral usage for each of eight coarse categories plotted for (a) Cntnap2 KO and (b) L7-Tsc1 mutant mice for each of four observation days. All individuals are shown as points, colored traces correspond to the median fraction of time spent in the behavior for each day. c) The usage of turning and locomotion behaviors during the 20-min observation period for Days 1 through 4 for Cntnap2 knockouts and littermates (left) and L7-Tsc1 mutants and littermates (right). Colors indicate the strain (blue-negative, green-heterozygote, orange-homozygote). Shaded regions represent the 95% confidence interval.

The defect in L7-Tsc1 mutant habituation was most apparent in locomotion and turning behaviors (Figs. 5 and S5). These mice did not show the expected reduction in turning either over days or within the 20-min observation period. Turning was used more often, but did not decrease to the same degree over the observation time as littermates. Locomotion decreased over time but to a lesser degree and the level of locomotion in the L7-Tsc1 mutant mice was always higher than in littermates.

### 2.5 Grooming behaviors vary in a complex manner in ASD models

Grooming refers to a variety of repetitive self-touching behaviors, and mouse self-grooming has been used as an animal model for the self-stimulating behaviors observed in autism [11]. Previous reports have limited quantification to 10 minutes of observation and have not distinguished different types of self-grooming or reported sex differences [42,46]. In our four-day recording period, we observed that all mouse groups groomed more frequently as the days passed (Figs. 4a, 5a,b, 6), with larger increases in males compared with females. We furthermore found that within-model contrasts (male v. female, Cntnap2 KO v. Cntnap2 wild type, L7-Tsc1 mutant v. L7-Tsc1 wild type) were modest on Day 1 but grew considerably over the next three days. The two ASD models showed opposite trends: Cntnap2 KO mice groomed more frequently than their littermates, whereas L7-Tsc1 mutants mice groomed much less frequently (Fig. 6).

**Figure 6:**
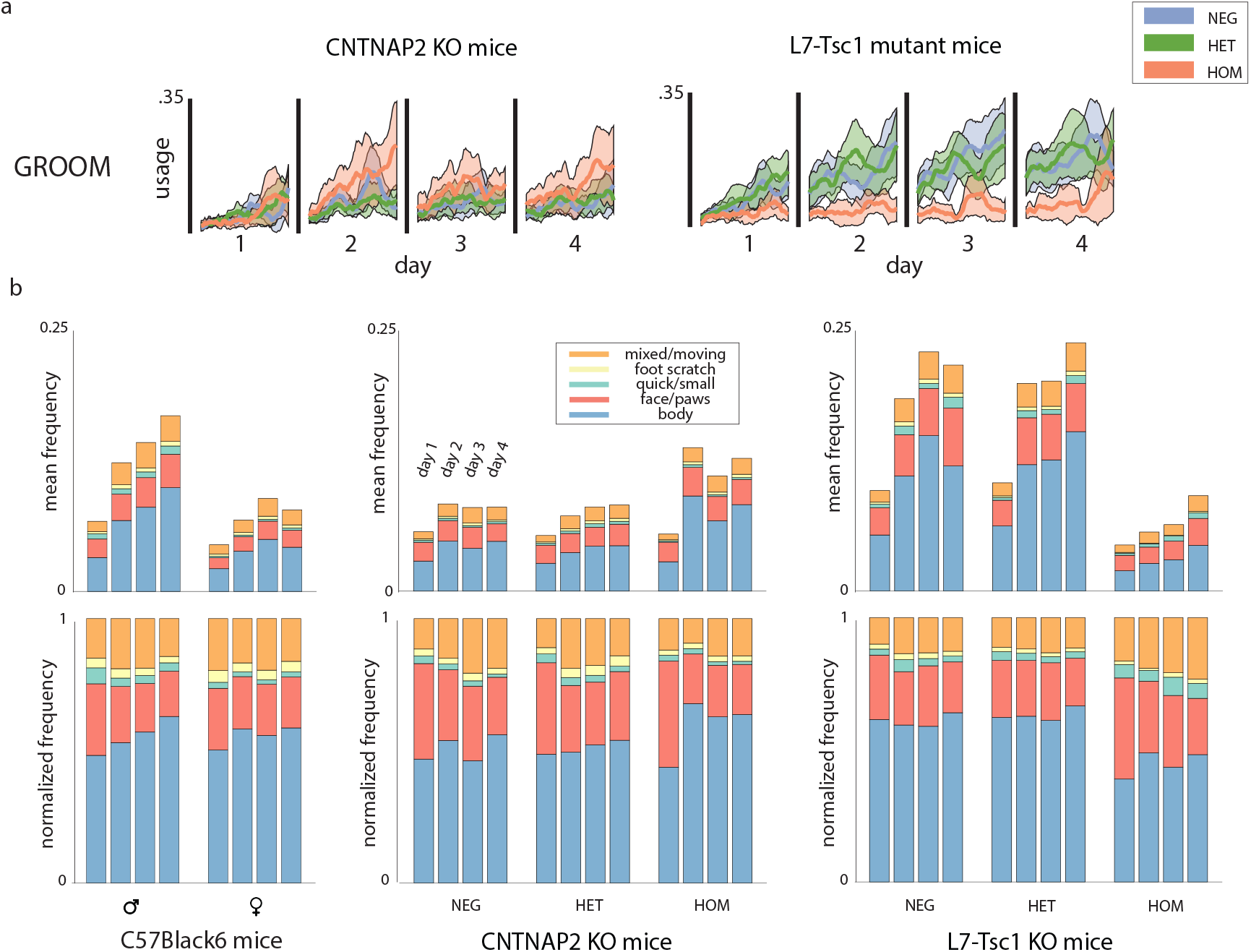
Habituation of grooming behaviors. a) The usage of grooming behaviors during the 20-min observation period for Days 1 through 4 for Cntnap2 and L7-Tsc1 mice. Colors indicate the strain (blue-negative, green-heterozygote, orange-homozygote). Shaded regions represent the 95% confidence interval. b) Stacked bar plots showing the mean frequency across mice for each of five grooming behaviors across Days 1 through 4 with corresponding stacked bar plots below showing the normalized mean frequency for each of five grooming behaviors.

The grooming class could be further broken up into several modes of grooming: body grooming, face/paw grooming/licking, small quick movements, foot scratching, and a mixture mode to account for time spent readjusting between modes. The total number of episodes spent grooming was normalized to 1 to obtain a relative usage of these grooming modes (Fig. 6c). Despite differences in total grooming frequency between C57BL/6J males and females, their relative grooming-mode frequencies were similar and showed the same trend over time. Across days, body grooming increased in relative frequency while face grooming and quick grooming movements became less frequent. Both ASD models showed altered distributions of the grooming behaviors. Cntnap2 KO mice exhibited more body grooming compared to their littermates on Days 2 through 4. L7-Tsc1 mutant mice showed the opposite effect with considerably less body grooming and more face/paw grooming than wild type littermates on all days of observation.

Grooming frequency also increased dramatically within each day’s observation period. Cntnap2 KO mice and their littermates groomed with similar frequency at the start of each day’s observation, but knockouts increased at a greater rate during each 20-minute period (Fig. 6a). Conversely, L7-Tsc1 wild-type and heterozygote mice groomed more as each day progressed, but mutant mice did not. These measurements show that systematic patterns of variation in grooming within each day of observation typically exceeded the variation across days.

### 2.6 L7-Tsc1 mutant and Cntnap2 KO mice show locomotion speed and gait defects

All experimental conditions showed reduced amounts of locomotion after Day 1. However, the total amount of locomotion and the distribution of locomotion speeds depended on condition. Male and female C57BL/6J mice showed a similar trend in locomotion amount and speed with the female mice moving on average 17% faster than their male counterparts (Fig. 7a). Cntnap2 KO mice did not differ from their littermates in the amount or speed of locomotion. In contrast, while L7-Tsc1 mutant mice locomoted more than either the C57BL/6J mice or their heterozygote littermates, they did so at a 15-25% slower speed. These differences persisted across all four days of testing.

**Figure 7:**
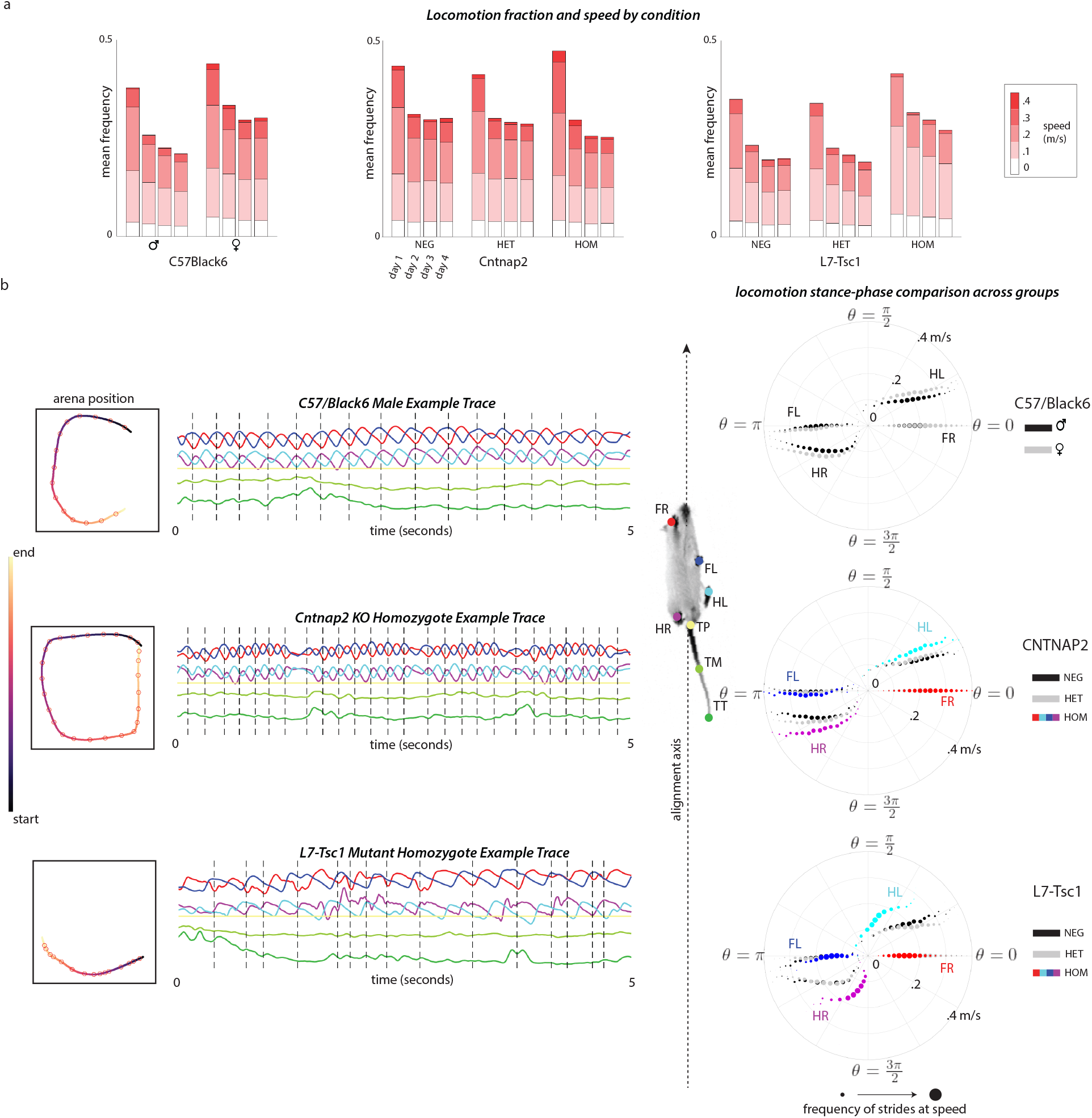
Locomotion kinematics are altered in the ASD models. a) Stacked bar plots showing the mean frequency for each of four speed bins on Days 1 through 4. b) Left: Examples of the motion of the limb and tail points during 5-second bouts of locomotion for C57BL/6J, Cntnap2 KO, and L7-Tsc1 mutants. The trajectory of the centroid of the mouse is color coded by time. Time series of the position of each body part projected onto the anterior-posterior axis is plotted. The start of each gait cycle is marked with a dashed black line and defined using the front right paw (FR, red). Center: An example image of a mouse is labeled with colors used for each body part, and the axis of body alignment used when segmenting strides is shown as the axis between the nose and tail base point (TP). Right: Polar plots of the phase *θ* of the gait cycle in which each paw reaches a minimum along the alignment axis in the body frame. The front right paw is used to define *θ* = 0. The radial axis indicates the speed of locomotion, which was used to bin the phase results, and the size of the circles indicate how frequent that particular speed was for a given condition. For Cntnap2 KO and L7-Tsc1 mutants, the colors of dots correspond to the four labeled paw points.

Nearly all children with autism show motor impairment [18,22], with gait following a locomotor pattern resembling cerebellar ataxia [16,37]. To explore the kinematics of locomotion, we identified all locomotion bouts (Figs. 1g and 7b) and analyzed individual body part trajectories along the anterior-posterior axis. The front and hind paws moved in an oscillating pattern in the frame of reference of the mouse as expected (Fig. 7b) [28]. We also found that the tail often oscillated as well, matching the frequency of the paw oscillation.

Mouse locomotion could be broken into two different gaits depending on the speed. At very slow speeds (*υ* < 0.1 m/s), C57BL/6J mice used a *walking* gait with each paw moving for approximately one quarter of the gait cycle (Fig. 7d). The order of movement and coordination in the walking gait was Front Right, Hind Left, Front Left, Hind Right. At higher speeds, mice transitioned to a *trot* gait with alternating pairs of legs moving together. However, the opposite pairs of legs are not in phase with each other within the gait cycle. We found that for speeds above ~ 0.1 m/s, C57BL/6J mice had an average phase difference of ~ 0.5 rad, or about 8 percent of the gait cycle. This phase offset decreased with speed from about ~ 1 rad at *υ* = 0.1 m/s to ~ 0.25 rad at *υ* = 0.4 m/s.

The ASD mutants displayed differences in both the gait transition speed and the phase difference between opposite limbs. Cntnap2 KO mice showed a similar transition speed of about 0.1 m/s, as did to their littermates. In contrast, L7-Tsc1 mutants used the walking gait much more and transitioned to the trotting gait at a much higher velocity of ~ 0.2 m/s. The phase difference during trotting was larger for both ASD models. Cntnap2 KO had a phase difference at 0.2 m/s of 0.8 rad compared to a difference of 0.6 rad for their littermates. L7-Tsc1 mutants had an even larger difference of 1.0 rad at 0.2 m/s compared to 0.5 rad for their littermates.

## 3 Discussion

We have found that the observation of mice in an open arena is sufficient to characterize behavior on scales ranging from sub-second individual limb movements to multi-day adaptive change. The two mouse models we used, Cntnap2 KO and L7-Tsc1 mutants, differed in distinct ways from wild-type mice on within-day measures of behavior, including gait defects and self-grooming. In addition, L7-Tsc1 mutant mice showed defects in multi-day behavioral evolution. Thus our methods can quantify complex behavior in ways that recall human autism, and differentiate between mouse models’ capacity to express these phenotypes (Fig. 8).

**Figure 8:**
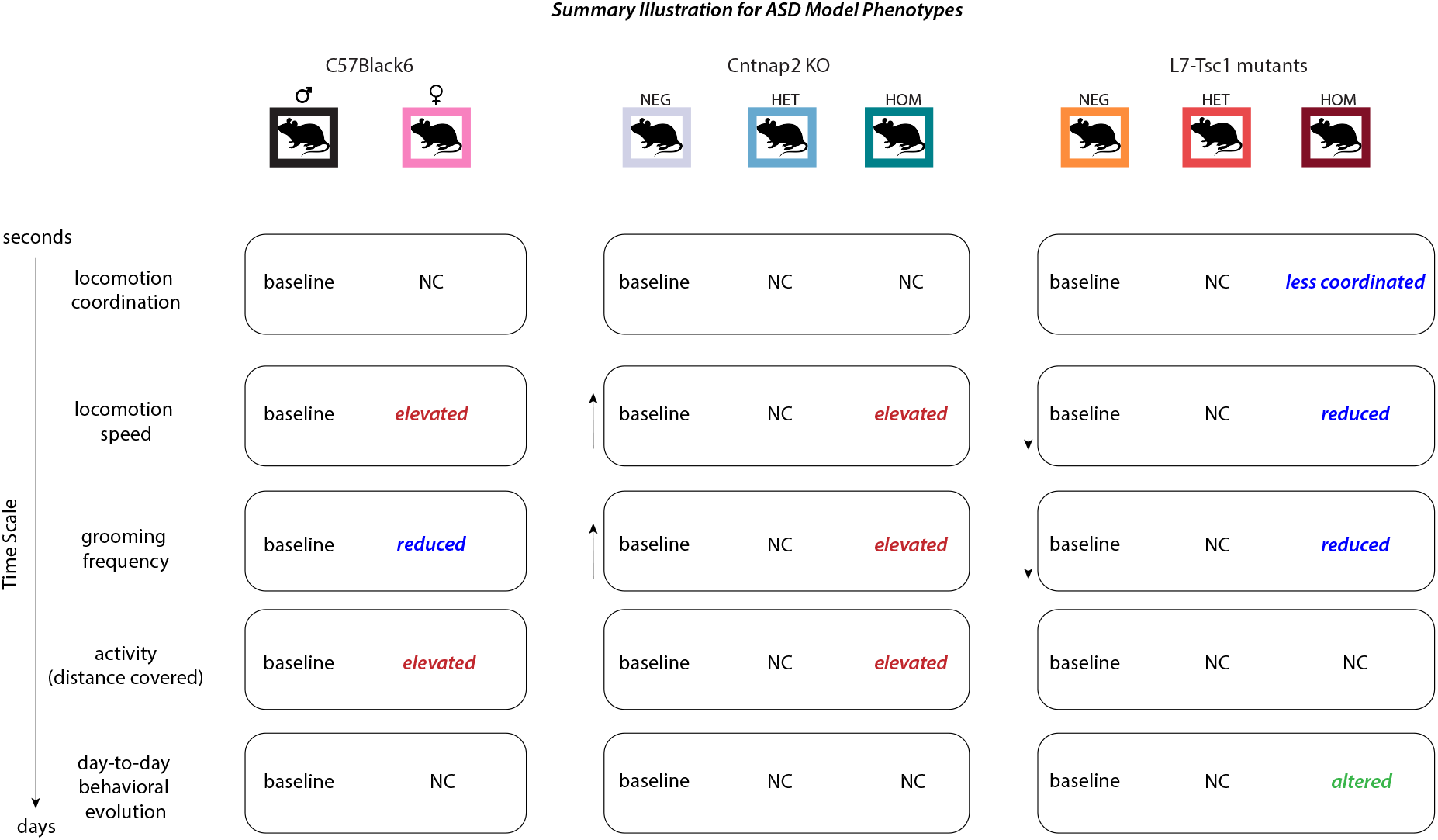
Summary of behavioral phenotypes of experimental groups

Animal behavior is high dimensional. Changes in movement, transitions between states, and adaptive change are most easily observed by different metrics on multiple time scales. Our method for semi-supervised behavioral classification allows exploration of all three types of dynamics from a single video recording. Such deep behavioral phenotyping can capture rich behavioral readouts that illuminate the effects of genetic and experimental perturbations in naturalistic environments.

Previous methods for analyzing freely-moving behavior have been limited to restricted feature sets. Observations from freely-moving animals have focused on simple measures, such as spatial location, using footprint measurement and centroid tracking. Detailed limb movements have been observed in constrained environments such as a corridor or treadmill to model the relationship between gait and postural features [28, 29], but this approach does not capture movement in a more natural context. Our approach allows all these types of measurements to be made at once in an automated manner.

The simplicity of the open field allows rapid characterization without need to train mouse or experimenter. This level of automation and standardization may contribute to differences with past work. For example, we found that L7-Tsc1 mutant mice showed reduced grooming compared with wild-types, yet past studies report increased grooming. We found that grooming included many sub-states that involved different movements. Differences in how human scorers classify such states may lead to lab-to-lab variability in quantification. Additional sources of variation include conditions that facilitate grooming, such as spraying mice with water or including bedding material in the chamber. Conversely, our automated methods incorporated a temporal window for clustering, so that singular quick grooming movements might have been missed. Further study is needed to reconcile these approaches.

Previous studies of mouse behavior have identified large sex differences in pairwise interactions (i.e. same-sex vs. opposite-sex interactions). Our analysis identified robust sex differences even in the absence of a specific task or another mouse in the chamber. The male-versus-female difference was also seen in Cntnap2 KO-versus-wild type mice, where knockouts tended to display increased activity. In contrast, L7-Tsc1 mutant mice showed a reversed phenotype from the Cntnap2 KO mice and presented with slower and less-coordinated behaviors. Our findings recall the observation that men are identified as being on the autism spectrum four times as often as women, and would be consistent with the possibility that regulation of some genes (by our measures, Cntnap2) but not others (Tsc1) might have differential effects by sex [31].

Mouse models of autism are typically generated from a focal perturbation affecting as little as one gene. However, a single-gene disruption can still affect many cell types and brain regions at once. Cntnap2 is expressed in neocortex, striatum, hippocampus, and cerebellum [32]. Therefore behavioral changes in Cntnap2 KO mice may arise from perturbation of any of these regions or a combination of them. In contrast, L7-Tsc1 mice exhibit reduced Tsc1 expression specifically in Purkinje cells, thus influencing behavior through cerebellar impacts on brain activity. The patterns of disruption are therefore likely to differ. For example, Cntnap2 KO mice show reductions in neocortical spine number [19] whereas L7-Tsc1 mutant mice show increased neocortical spine density [23].

The ability to simultaneously characterize movement kinematics and larger scale features of behavior, even without a task condition, has particular relevance to the study of autism. Movement disruptions appear in 80 percent of children with autism [18, 22], despite the fact that movement is not considered part of the classic triad of defining symptoms. Injury to the cerebellum at birth leads to a sharply increased likelihood of autism, raising the possibility that the same structure that regulates smooth movement may also play a key role in driving the development of higher-level behavioral capacity [50]). In this way, movement and cognitive maturation may use shared neural substrates. Our analysis of Cntnap2 KO and L7-Tsc1 mutant mice reveals a shared disruption to specific parameters of gait arising from different genetic perturbations, one brain-wide and one cerebellar. We find that each model exemplifies different features of ASD studied in humans: hyperactivity and perseveration in one case, and failure-to-adapt and gait defects in another. In the future it may be possible to use alterations in movement to aid in the classification of autistic individuals. In addition, early-life identification of motor symptoms may allow rapid risk stratification for early intervention.

Mouse tests for autism-like phenotypes are selected for their putative relevance to human behavior and for their ease of administration. However, it is often difficult to probe systems-level mechanisms responsible for the observed alterations in performance. For example, Cntnap2 KO mice [8, 32] show disruptions on a wide array of tests, none of which are well studied at a circuit level. Deep behavioral phenotyping allows systems-specific defects (gait) and higher-order performance to be assayed within the same behavioral recording. In this way our methods open the future possibility of investigating complex behavior with substantially increased depth, both of phenotyping and of neural mechanism. The power of these methods will increase as they are integrated with specific task or test conditions, as well as neural recording and perturbation.

Our approach to characterizing behavior may also eventually be useful in the understanding of human autism. Predictors of autism such as unusual patterns of sensory response are known to emerge in the first year of life. Deep behavioral phenotyping can expand the range of observations without need for a specific task. By characterizing fine-grained movement, limb coordination, and long-time-scale adaptation, it may be possible to identify distinctive features of behavior that precede a formal diagnosis of ASD. Such information can potentially stratify at-risk children according to different points of departure from neurotypical paths of cognitive and social development. As the brain-wide mechanisms of autism also become better understood, such stratification may aid in the understanding and treatment of the many forms of autism.

## 4 Methods

### 4.1 Experimental animals

C57BL/6J male (*n* = 60) and female (*n* = 20) were ordered from Jackson Laboratory (The Jackson Laboratory, Bar Harbor, ME) and had at least 48 hours of acclimation in the Princeton Neuroscience Institute vivarium before experimental procedures. Two mouse models were used to analyze autistic endophenotypes.

L7-Tsc1: To test cerebellar modulation of naturalistic behaviors, a Purkinje cell degeneration model with a tuberous sclerosis 1 gene mutation was used [26,46]; L7Cre;Tsc1flox/flox). Initially, Tsc1flox/flox (Tsc1tm1Djk/J, Jackson Laboratory stock #005680) mutant mice were crossed into L7/Pcp2 mice (B6.129-Tg(Pcp2-cre)2Mpin/J, Jackson Laboratory stock 004146) to create a Purkinje cell specific mutation L7Cre;Tsc1 flox/+. The progeny used from this cross are control (L7Cre;Tsc1+/+), heterozygous (L7Cre;Tsc1flox/+) mice and mutant (L7Cre;Tsc1flox/flox) mice. Only male animals were used for behavior experiments. Mice are of mixed genetic backgrounds (C57Bl/6J, 129 SvJae and BALB/cJ).

Cntnap2: A knockout of Cntnap2 associated with cortical dysplasia-focal epilepsy, Cntnap2-/- (B6.129(Cg)-Cntnap2tm1Pele/J, Jackson Laboratory stock 017482) was bred to C57BL/6J (Jackson Laboratory stock #000664) male mice to obtain heterozygote (Cntnap2+/-) progeny. These litters were then bred as a heterozygote strategy to obtain litters with wild type (Cntnap2+/+), full KO (Cntnap2-/-), and heterozygote (Cntnap2+/-) mice.

All animals were tested in adulthood (>10 weeks of age) and housed with four littermates per cage in Optimice cages (Animal Care Systems, Centennial, CO). PicoLab Rodent Diet food pellets (LabDiet, St. Louis, MO) and drinking water were provided ad libitum. All mice were kept in a reverse light cycle for behavioral testing. All experimental procedures were approved by the Princeton University Institutional Animal Care and Use Committee and performed in accordance with animal welfare of National Institutes of Health.

### 4.2 Open Field Test recordings

Animals were placed in an open field area measuring 45.72 cm^2^ and 30.48 cm in height with a transparent polycarbonate floor. A Point Grey grayscale camera (12-bit grayscale, 1280 × 1024 pixel resolution at 80 HZ) was used to image animals from below. The ventilated soundproof box was illuminated with far-red LEDs. To prevent noise disturbance, doors were kept closed during acquisition. Each mouse was recorded for 20 minutes before being returned to group housing. Movies were saved as h5 files for further processing in MATLAB.

### 4.3 Centroid tracking and clipping

Full-frame movies were imaged at 1280 × 1024 pixel resolution, and contain the entire arena in their field of view. Mouse centroids were tracked and movies are clipped prior to performing joint tracking. Each movie was first sampled at random to generate a median image from 50 frames across the experiment. 1000-frame chunks were then loaded and each frame was median-subtracted, down-sampled, and Gaussian-blurred. The centroid of the brightest point after this procedure identified the approximate location of the centroid of the mouse, and a 400 × 400 pixel frame centered at these coordinates was kept in memory after mean-subtraction. Finally the fully-resolved clipped movie was saved as an h5 file for further analysis, along with an information file containing the coordinates used to clip the movie from the original frame. As a validation of comparability with past work, center-of-mass tracking [39] showed that in all experimental groups, mice covered the most distance on their first day of exposure to the open field, declining on later days [6,38].

### 4.4 Frame alignment and preprocessing

350 frames sampled from among clipped movies were used to train a deep neural network (DNN) using LEAP in order to identify the coordinates of the snout, center point, and tail point (where the tail meets the body) of the mouse [33]. Each movie was then labeled using this network. The resulting body part coordinates were loaded along with each clipped movie and info file, and each frame of the original clipped movie was centered at the tail point and aligned to the tail point-to-nose axis using image translation and rotation functions in MATLAB. The centroid information was updated to incorporate the new centroid after translation, and the rotation values were also saved for each frame in an updated information .mat file.

### 4.5 Body-part coordinate identification

We used 660 frames to train a deep neural network (DNN) from the aligned clipped movies described in the previous methods [33]. The neural network was trained iteratively using the LEAP interface, where the network was used to label a sample set of frames that were then fixed manually and used as training data in order to update the network. In particular, the training set was designed to include examples of various postures during diverse behaviors that the network failed to label in early iterations. Frames with multiple occlusions of the head and limbs during grooming were trained with a ‘best guess’ of where the body part in question would be located, and the network was updated until all postures were labeled satisfactorily. Overall the network was trained to identify 18 body points (snout, ears, chin, inner and outer limbs, center, sides, tail point, tail center, and tail tip), which would be used for behavioral classification.

### 4.6 Classical measures using centroid tracking

Mouse centroid coordinates throughout each open field recording were used to calculate basic measures of activity and spatial occupancy for each experimental trial. The centroid was used to calculate the velocity and position of the animal at any given time-point, and this value was further used to calculate the spatial distribution of animals in the arena and a number of other metrics such as center-crossings throughout all recordings.

### 4.7 Postural representations over time

Mouse bodies are not rigid and the choice of axis for egocentric alignment introduces bias that depends on the relative position of body parts used to establish alignment. Instead of aligning to a body axis, we used the distances between all pairs of virtual marker coordinates to capture the posture of the animal at each frame. The distance matrix for a given frame *D_n_* was calculated as follows.

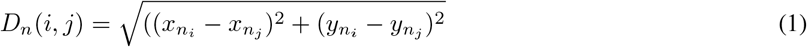

The posture of an animal over a recording is now represented as an 11 × 11 × *n* array where *n* is the number of frames in a given movie. We performed online PCA on the entire set of movies by sampling batches of frames and calculating the mean and covariance of *D* for each batch, keeping a running tab of the total mean and covariance across all movies and finally finding the eigenvectors of the ultimate covariance matrix. We projected the distance matrix onto the top 10 of these eigenvectors to produce a set of projections that captured about 90 percent of the variance in *D* with a greatly reduced dimensionality. The 10 × *n* array describing the projections along the top ten principal components was saved for each movie and captured the posture of each animal over time.

### 4.8 Temporal representation of behavior

We performed a wavelet decomposition on the projection matrix containing the top ten posture modes in order to find the power spectrum across a dyadically spaced set of frequencies between .25Hz and 20Hz. The 10 × *n* postural representation was transformed into a (10 × 25) × *n* wavelet spectrogram where each of the ten postural modes was described by power along 25 frequencies. The relative power among different frequencies corresponds to how quickly the posture of the animal is changing in time, and power in high frequencies means that the projection along a particular postural mode is changing quickly. The postural-dynamic representation described by the wavelets was the final signal used to compare time points from behavioral movies. We used the log of this signal, with all values below −3 set to −3 to cluster all recorded behavior into fine- and coarse-grained categories that were used in all further behavioral descriptions. The same transformation was applied to the wavelets from each movie during the reembedding step.

### 4.9 Clustering and dimensionality reduction to form behavioral classes

Because of the unbalanced nature of our data, where extremely specific yet common behaviors like locomotion and a particular speed might overwhelm an automated clustering algorithm, we first used importance sampling to design a representative behavioral sample set, like we did with postures for training the DNN. We sampled movies from each experimental condition and created a dataset that encompassed all of the types of movements that were found in our recordings. This was particularly important for labeling uncommon behaviors that may not have appeared in many movies but were still behaviorally interesting.

In order to sample all behaviors present among our recordings, we first performed dimensionality reduction on the time series *x_i_* from each movie independently using tSNE [47]. The resulting maps were sampled to find a variety of templates, or examples of the wavelet values, that were present in the given movie, ensuring that even rare templates were represented. These templates, on the order of 100 per movie used, were concatenated to form a training set that was then clustered into classes that formed the basis of our behavioral labels. We performed clustering using two methods that are complimentary when visualizing data: tSNE mapping and *k*-means clustering. First, as previously described [5], we performed dimensionality reduction on a training set of ~ 40,000 samples by embedding the 250-dimensional wavelet signal into a 2-dimensional tSNE map. The final map can be used to cluster behaviors using the watershed transformation, or to visualize the density of the behavioral repertoire used by an animal or group of animals. Instead of clustering in the 2-dimensional map, we also performed *k*-means clustering on the training set, with a target of k=100 clusters. Comparing the tSNE and *k*-means methods reveals similar results, however it is simpler to embed new samples into the *k*-means clusters by simply finding the nearest neighbor of a given sample, and assigning it the cluster designation of the nearest neighbor.

### 4.10 Posture-dynamical fingerprinting

The power in the wavelet spectrum across tracked body parts could be used to interpret the results of behavioral clustering. For interpretability, the wavelet decomposition was recalculated using the raw displacement in real space for each tracked body part, in lieu of performing PCA to find a postural representation. The body part wavelet signals associated with time-points assigned each to a given behavioral class were averaged and reshaped to generate a power spectrum. These ‘fingerprints’ detail the power in each frequency used in the decomposition for each tracked body part, and correspond to visible features of the behaviors found in each cluster or class.

### 4.11 Calculating behavioral time-series and normalized usage of behavioral classes

Each 20-minute behavioral movie contained approximately 96,000 frames sampled at 80 Hz. The posture-dynamics clustering method we described previously was used to assign a behavioral label or class, from 1 to 100, to each of these frames. The creation of 100 clusters, more than were easily distinguishable by the human eye, allowed us to manually curate behavioral groups by viewing randomly-sampled examples from each cluster. We then grouped behaviors into coarse groups that aligned with an understanding of behavior and the rodent phenotyping literature, and also were already combined into building blocks based on our automated method.

Usage during a given experiment or segment of experiment was defined as the normalized histogram over behavioral classes. Evolution of behavior over the course of an experiment was found by choosing a time-window, for example 180-seconds, and calculating this histogram over a sliding window of this size over the course of the experiment. The 95% confidence interval was calculated using the density values from each individual mouse.

### 4.12 Finding locomotion bouts, steps, and phase

Locomotion was first identified using the *k*-means clustering method. All time-points that were identified as belonging to the locomotion class were found, and a structure of joint trajectories and centroid positions of bouts over 500 ms long was created for each movie. Each locomotion bout was then further characterized. First all bouts were up-sampled by a factor of 100 and *Z*-scored, and the findpeaks function in MATLAB was used to locate the time indices of the beginning of each stride for each of the four limbs. Peaks were required to be at least 80 ms apart and with a minimum peak prominence of .2 after *Z*-scoring. The peaks were used to assign a phase to each limb at each time-point, where the end of the swing, or upward motion of the paw, corresponds to *θ* = 0. For each time at which the front right paw has a phase of *θ* = 0, the phase of each other limb was sampled and recorded, along with the centroid speed and angular velocity of the animal, and the arena coordinates at which the step was performed.

## 5 Data availability statement

All data required to reproduce results are available at Princeton Library Research Databases. All code for the mouse LEAP models and accompanying metrics are packed with skeleton and labeling data. Raw behavior movies are available upon request. Database with time-series of predicted positions of virtual markers, training dataset, and PCA components can be downloaded at digital DataSpace repository (https://doi.org/10.34770/bzkz-j672). The Github repository (https://github.com/PrincetonUniversity/MouseMotionMapper) is set up to use the batch processing system Slurm and contains the code to reproduce the analysis pipeline. The repository contains code for reproducing all figures in the main text and supplement. Examples of preprocessing steps, unsupervised classification, reembedding, fingerprinting, and all other utility functions used are also included.

## 6 Acknowledgments

We thank Silke Bergeler and Gordon Berman for discussion and comments on an earlier version of the manuscript; and Laura Lynch, Thomas Kellogg and Dafina Pacuku for technical assistance. This work was supported by NIH R01 NS045193, R01 MH115750, and U19 NS104648.

## 7 Author Contributions

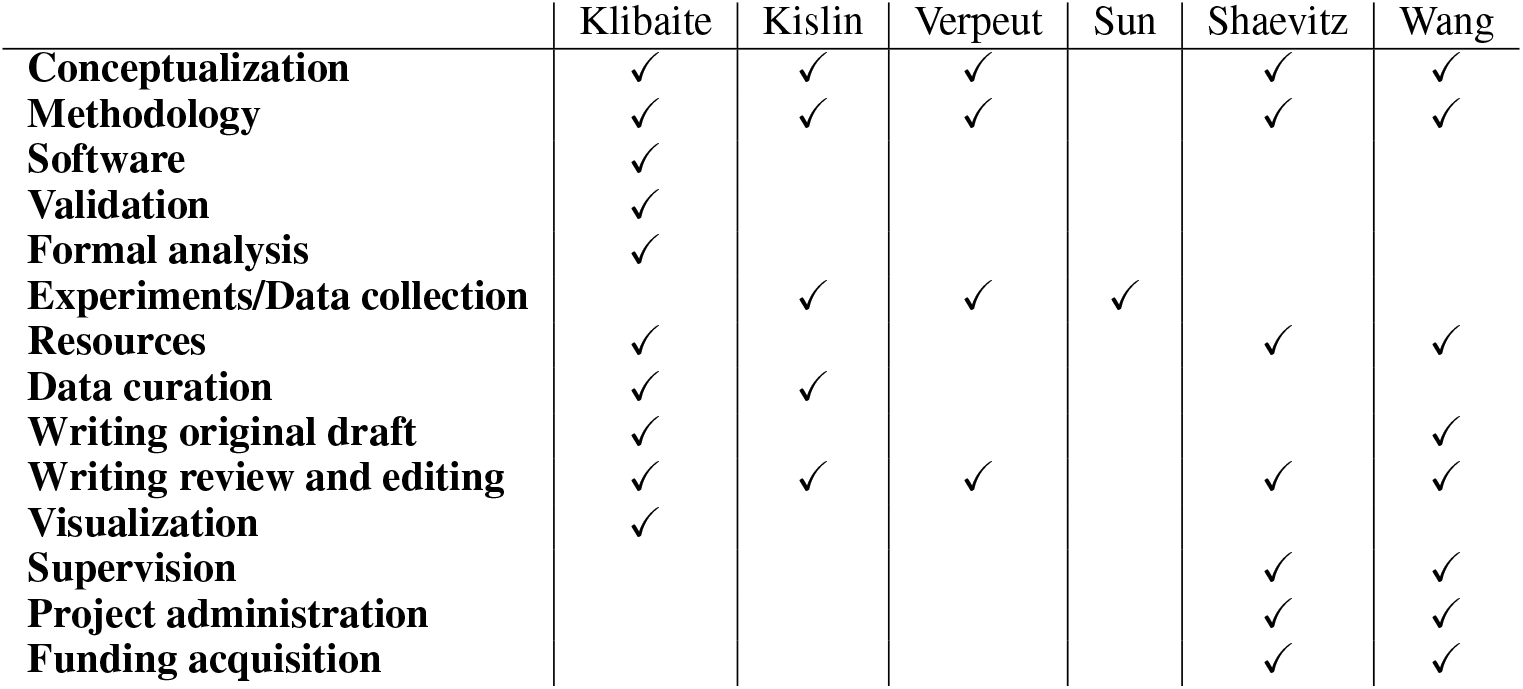

## Supplementary Materials

**Figure S1: Supplement to Figure 1.**
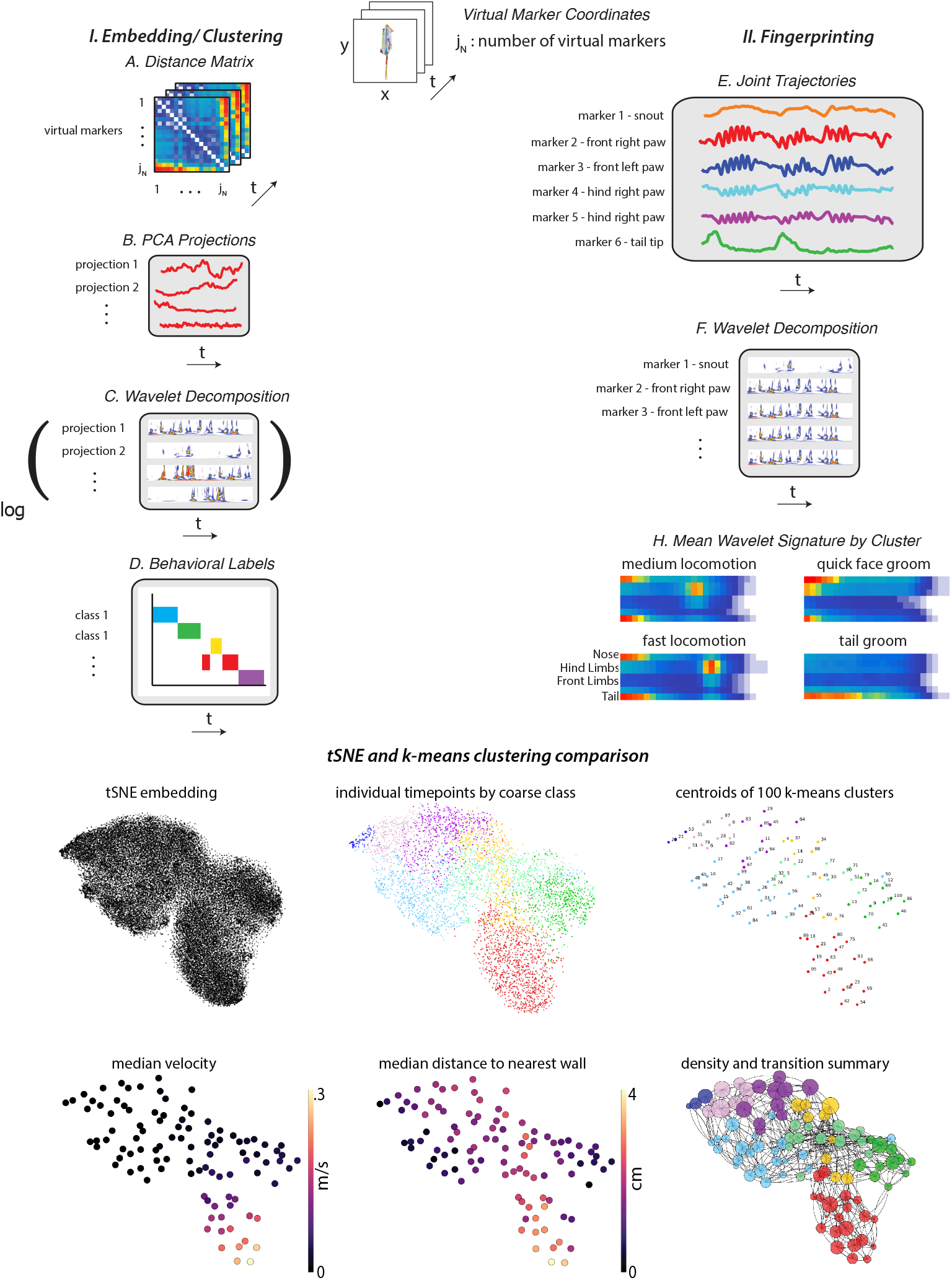
I. A visual representation of the calculations performed to find the 100 *k*-means clustered classes is described by steps A-D. II. Raw joint trajectories are used to create wavelet signatures or ‘behavioral fingerprints’ by finding the mean power spectrum during each behavioral class found in part I. Visualizations of the tSNE embedding for a sub-selection of data show how the methods are related.

**Figure S2: Supplement to Figure 3.**
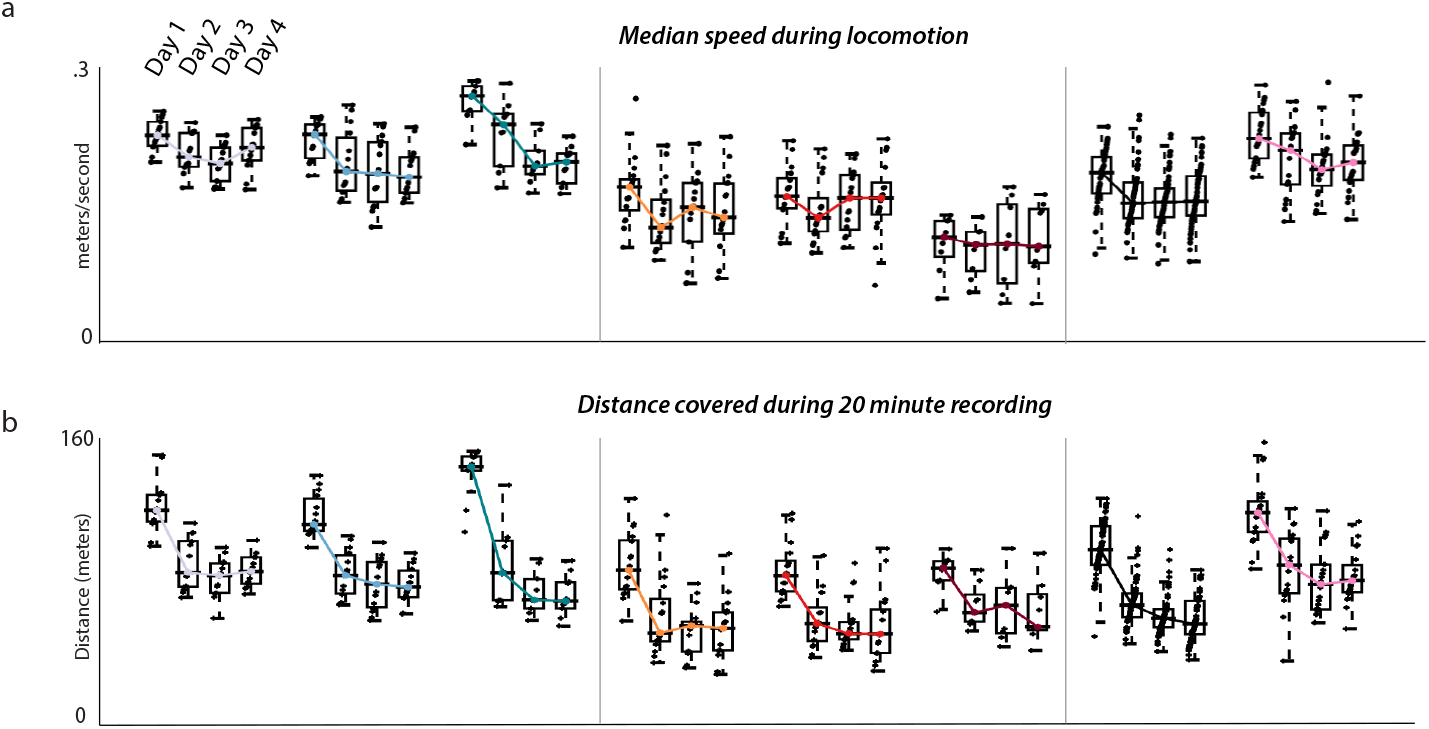
Common metrics for open field performance plotted for all conditions. A) Median velocity during locomotion, B) Median distance covered during 20 minutes in the open field.

**Figure S3:**
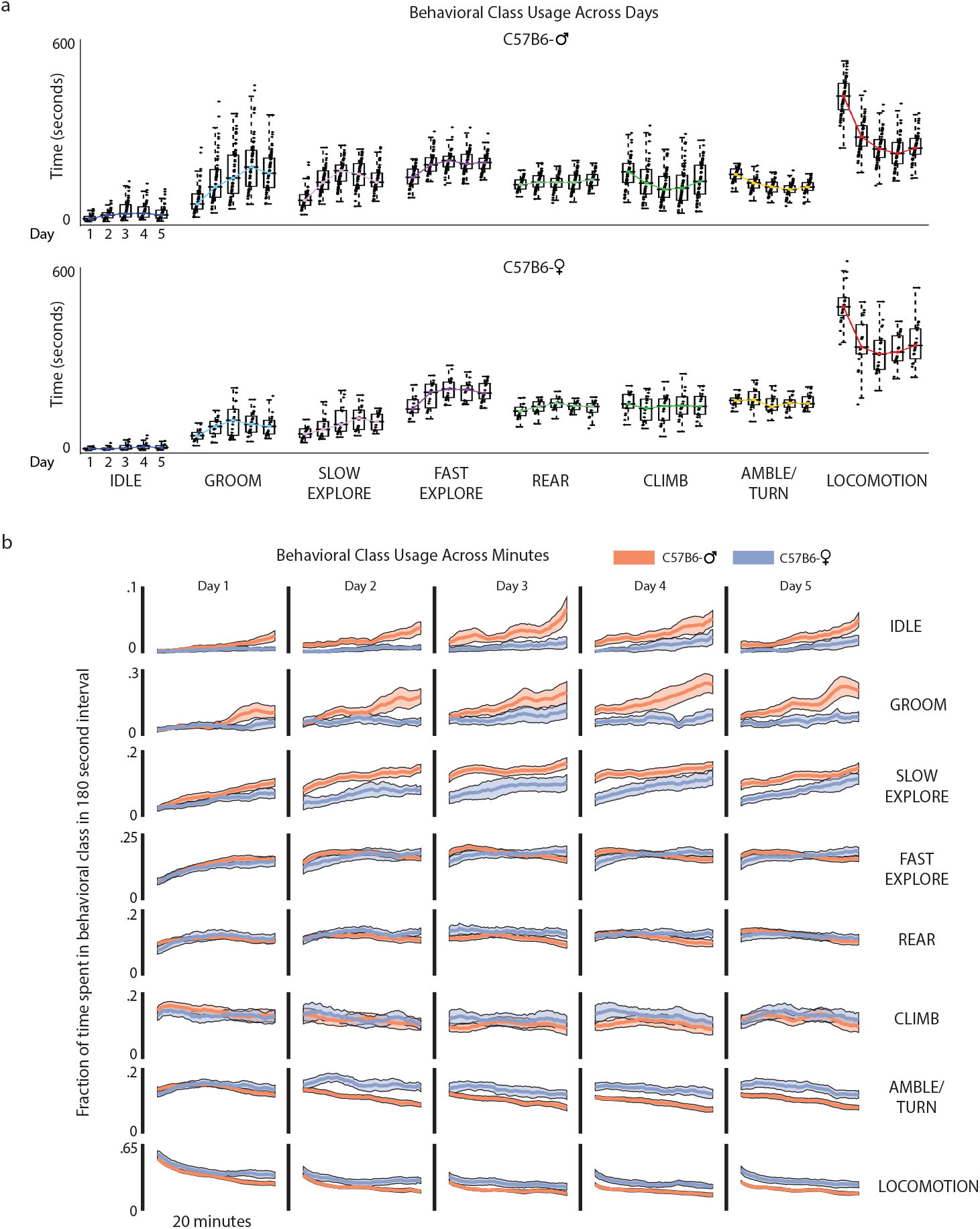
Behavioral summary of C57BL/6J male and female mice. a) Behavioral usage for each of eight coarse categories plotted for C57BL/6J male(top) and female (bottom) mice for each of five observation days. All individuals are shown as points, colored traces correspond to the median fraction of time spent in the behavior for each day. b) The mean usage of each coarse behavioral class during 20 minutes of observation for each of five days. Shaded regions represent 95% confidence interval.

**Figure S4:**
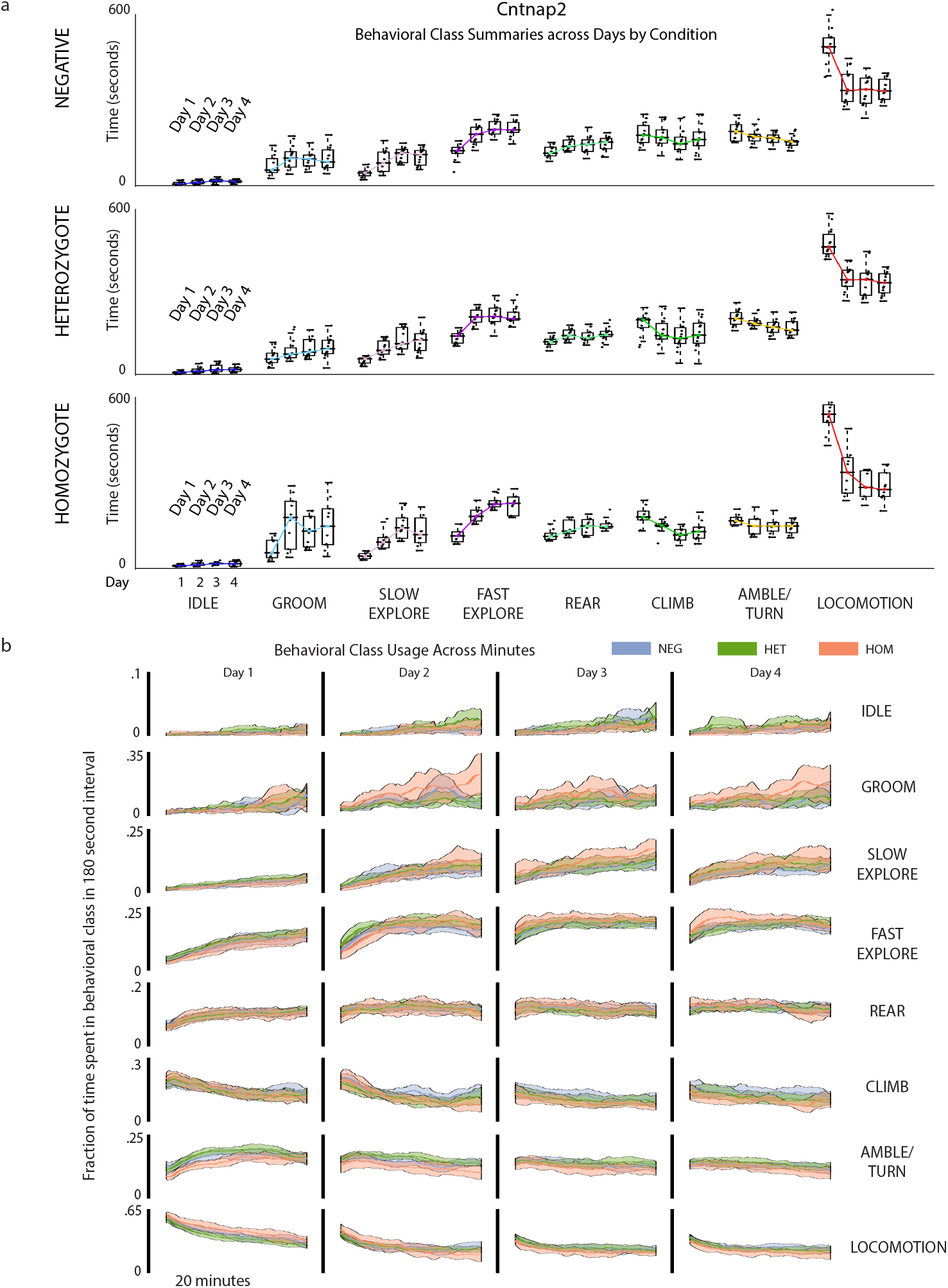
Behavioral summary of Cntnap2 KO mice. a) Behavioral usage for each of eight coarse categories plotted for Cntnap2 KO negative (top), heterozygote (middle), and homozygote (bottom) mice for each of four observation days. All individuals are shown as points, colored traces correspond to the median fraction of time spent in the behavior for each day. b) The mean usage of each coarse behavioral class during 20 minutes of observation for each of four days. Shaded regions represent 95% confidence interval.

**Figure S5:**
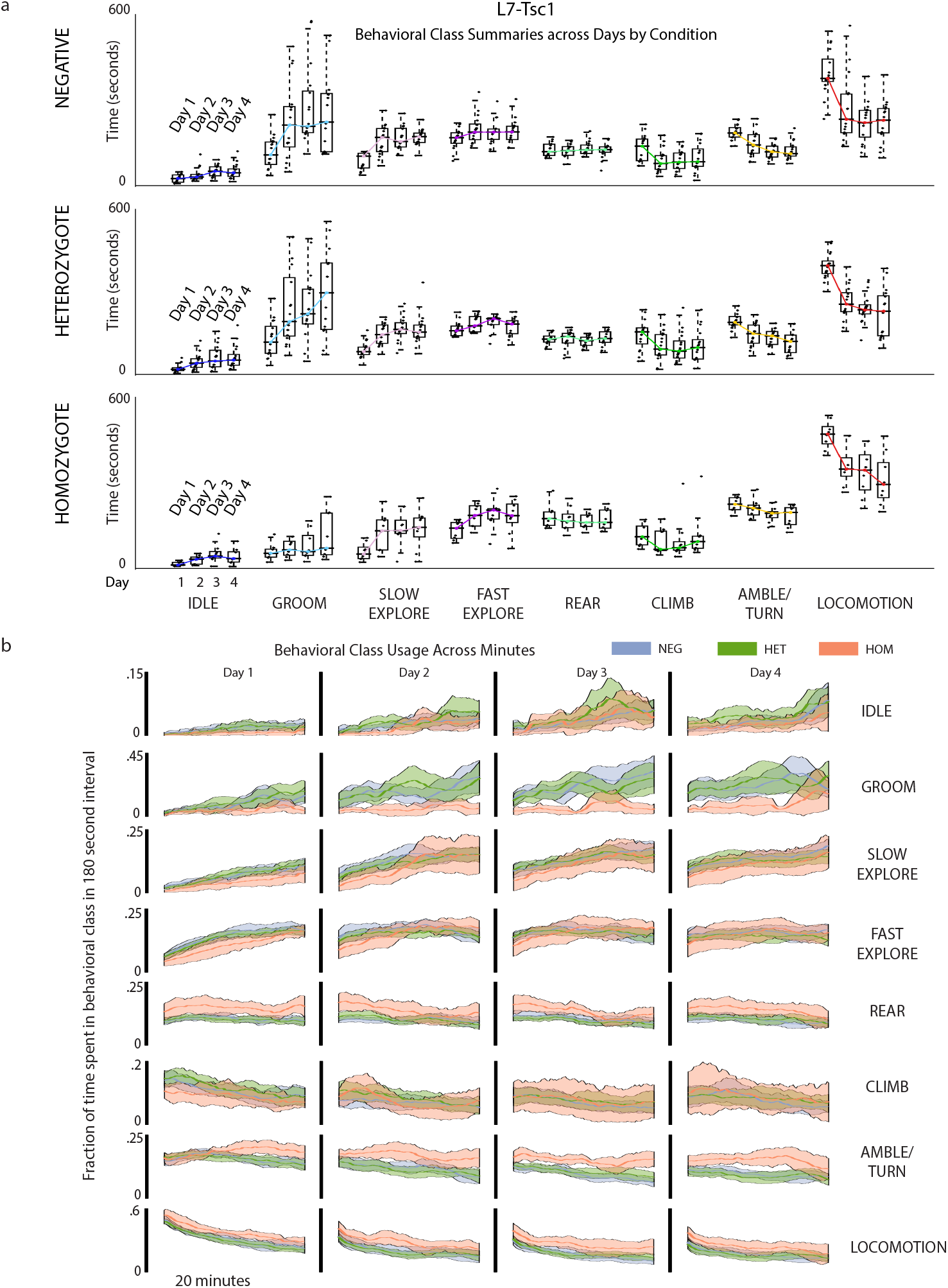
Behavioral summary of L7-Tsc1 mutant mice. a) Behavioral usage for each of eight coarse categories plotted for L7-Tsc1 mutant negative (top), heterozygote (middle), and homozygote (bottom) mice for each of four observation days. All individuals are shown as points, colored traces correspond to the median fraction of time spent in the behavior for each day. b) The mean usage of each coarse behavioral class during 20 minutes of observation for each of four days. Shaded regions represent 95% confidence interval.

## Notes

https://doi.org/10.34770/bzkz-j672

https://github.com/PrincetonUniversity/MouseMotionMapper

### Competing Interest Statement

The authors have declared no competing interest.

## References

[1] American Psychiatric Association et al. Diagnostic and statistical manual of mental disorders (DSM-5^®^). American Psychiatric Pub, 2013.

[2] Yoav Benjamini, Ehud Fonio, Tal Galili, Gregor Z Havkin, and Ilan Golani. Quantifying the buildup in extent and complexity of free exploration in mice. Proceedings of the National Academy of Sciences, 108(Supplement 3):15580–15587, 2011.

[3] Gordon J Berman. Measuring behavior across scales. BMC biology, 16(1):23, 2018.

[4] Gordon J Berman, William Bialek, and Joshua W Shaevitz. Predictability and hierarchy in drosophila behavior. Proceedings of the National Academy of Sciences, 113(42):11943–11948, 2016.

[5] Gordon J Berman, Daniel M Choi, William Bialek, and Joshua W Shaevitz. Mapping the stereotyped behaviour of freely moving fruit flies. Journal of The Royal Society Interface, 11(99):20140672, 2014.

[6] Carina Bodden, Sophie Siestrup, Rupert Palme, Sylvia Kaiser, Norbert Sachser, and S Helene Richter. Evidencebased severity assessment: Impact of repeated versus single open-field testing on welfare in c57bl/6j mice. Behavioural brain research, 336:261–268, 2018.

[7] André EX Brown and Benjamin De Bivort. Ethology as a physical science. Nature Physics, 14(7):653, 2018.

[8] Daniela Brunner, Patricia Kabitzke, Dansha He, Kimberly Cox, Lucinda Thiede, Taleen Hanania, Emily Sabath, Vadim Alexandrov, Michael Saxe, Elior Peles, et al. Comprehensive analysis of the 16p11. 2 deletion and null cntnap2 mouse models of autism spectrum disorder. PloS one, 2015.

[9] Marco Cambiaghi, Marco Cursi, Laura Magri, Valerio Castoldi, Giancarlo Comi, Fabio Minicucci, Rossella Galli, and Letizia Leocani. Behavioural and eeg effects of chronic rapamycin treatment in a mouse model of tuberous sclerosis complex. Neuropharmacology, 67:1–7, 2013.

[10] Kathryn K Chadman, Mu Yang, and Jacqueline N Crawley. Criteria for validating mouse models of psychiatric diseases. American Journal of Medical Genetics Part B: Neuropsychiatric Genetics, 150(1):1–11, 2009.

[11] Andrew D Chang, Victoria A Berges, Sunho J Chung, Gene Y Fridman, Jay M Baraban, and Irving M Reti. High-frequency stimulation at the subthalamic nucleus suppresses excessive self-grooming in autism-like mouse models. Neuropsychopharmacology, 41(7):1813–1821, 2016.

[12] Jacqueline N Crawley. Behavioral phenotyping of rodents. Comparative medicine, 53(2):140–146, 2003.

[13] Jacqueline N Crawley. Translational animal models of autism and neurodevelopmental disorders. Dialogues in clinical neuroscience, 14(3):293, 2012.

[14] Jacqueline N Crawley, John K Belknap, Allan Collins, John C Crabbe, Wayne Frankel, Norman Henderson, Robert J Hitzemann, Stephen C Maxson, Lucinda L Miner, Alcino J Silva, et al. Behavioral phenotypes of inbred mouse strains: implications and recommendations for molecular studies. Psychopharmacology, 132(2):107–124, 1997.

[15] Sandeep Robert Datta, David J Anderson, Kristin Branson, Pietro Perona, and Andrew Leifer. Computational neuroethology: A call to action. Neuron, 104(1):11–24, 2019.

[16] Gianluca Esposito and Paola Venuti. Analysis of toddlers’ gait after six months of independent walking to identify autism: a preliminary study. Perceptual and motor skills, 106(1):259–269, 2008.

[17] Sarah L Ferri, Ted Abel, and Edward S Brodkin. Sex differences in autism spectrum disorder: a review. Current psychiatry reports, 20(2):1–17, 2018.

[18] Kimberly A Fournier, Chris J Hass, Sagar K Naik, Neha Lodha, and James H Cauraugh. Motor coordination in autism spectrum disorders: a synthesis and meta-analysis. Journal of autism and developmental disorders, 40(10):1227–1240, 2010.

[19] Amos Gdalyahu, Maria Lazaro, Olga Penagarikano, Peyman Golshani, Joshua T Trachtenberg, and Daniel H Geschwind. The autism related protein contactin-associated protein-like 2 (cntnap2) stabilizes new spines: an in vivo mouse study. PloS one, 10(5):e0125633, 2015.

[20] Alex Gomez-Marin, Joseph J Paton, Adam R Kampff, Rui M Costa, and Zachary F Mainen. Big behavioral data: psychology, ethology and the foundations of neuroscience. Nature neuroscience, 17(11):1455, 2014.

[21] Todd D Gould. Mood and anxiety related phenotypes in mice: characterization using behavioral tests, volume 2. Springer, 2009.

[22] Dido Green, Tony Charman, Andrew Pickles, Susie Chandler, TOM Loucas, Emily Simonoff, and Gillian Baird. Impairment in movement skills of children with autistic spectrum disorders. Developmental Medicine & Child Neurology, 51(4):311–316, 2009.

[23] Verpeut J.L., Pisano T, Klibaite U., Kislin M., Lee J., Willmore L., Matl C., Pacuku D., Pereira T.D. and, Badura A.M., and Wang S.S.-H. Regulation of flexible learning, social interaction, and whole-brain cellular activity by lobule vi of posterior vermis.

[24] Ugne Klibaite, Gordon J Berman, Jessica Cande, David L Stern, and Joshua W Shaevitz. An unsupervised method for quantifying the behavior of paired animals. Physical biology, 14(1):015006, 2017.

[25] Ugne Klibaite and Joshua W Shaevitz. Paired fruit flies synchronize behavior: Uncovering social interactions in drosophila melanogaster. PLOS Computational Biology, 16(10):e1008230, 2020.

[26] David J Kwiatkowski, Hongbing Zhang, Jennifer L Bandura, Kristina M Heiberger, Michael Glogauer, Nisreen el Hashemite, and Hiroaki Onda. A mouse model of tsc1 reveals sex-dependent lethality from liver hemangiomas, and up-regulation of p70s6 kinase activity in tsc1 null cells. Human molecular genetics, 11(5):525–534, 2002.

[27] R Lathe. The individuality of mice. Genes, Brain and Behavior, 3(6):317–327, 2004.

[28] Ana S Machado, Dana M Darmohray, Joao Fayad, Hugo G Marques, and Megan R Carey. A quantitative framework for whole-body coordination reveals specific deficits in freely walking ataxic mice. Elife, 4:e07892, 2015.

[29] Ana S Machado, Hugo G Marques, Diogo F Duarte, Dana M Darmohray, and Megan R Carey. Shared and specific signatures of locomotor ataxia in mutant mice. bioRxiv, 2020.

[30] Eric J Nestler and Steven E Hyman. Animal models of neuropsychiatric disorders. Nature neuroscience, 13(10):1161, 2010.

[31] Meritxell Oliva, Manuel Muñoz-Aguirre, Sarah Kim-Hellmuth, Valentin Wucher, Ariel DH Gewirtz, Daniel J Cotter, Princy Parsana, Silva Kasela, Brunilda Balliu, Ana Viñuela, et al. The impact of sex on gene expression across human tissues. Science, 369(6509), 2020.

[32] Olga Peñagarikano, Brett S Abrahams, Edward I Herman, Kellen D Winden, Amos Gdalyahu, Hongmei Dong, Lisa I Sonnenblick, Robin Gruver, Joel Almajano, Anatol Bragin, et al. Absence of cntnap2 leads to epilepsy, neuronal migration abnormalities, and core autism-related deficits. Cell, 147(1):235–246, 2011.

[33] Talmo D Pereira, Diego E Aldarondo, Lindsay Willmore, Mikhail Kislin, Samuel S-H Wang, Mala Murthy, and Joshua W Shaevitz. Fast animal pose estimation using deep neural networks. Nature methods, 16(1):117, 2019.

[34] Talmo D Pereira, Joshua W Shaevitz, and Mala Murthy. Quantifying behavior to understand the brain. Nature Neuroscience, pages 1–13, 2020.

[35] Jan P Piek and Murray J Dyck. Sensory-motor deficits in children with developmental coordination disorder, attention deficit hyperactivity disorder and autistic disorder. Human movement science, 23(3-4):475–488, 2004.

[36] Nicole J Rinehart, Mark A Bellgrove, Bruce J Tonge, Avril V Brereton, Debra Howells-Rankin, and John L Bradshaw. An examination of movement kinematics in young people with high-functioning autism and asperger’s disorder: further evidence for a motor planning deficit. Journal of autism and developmental disorders, 36(6):757–767, 2006.

[37] Nicole J Rinehart, Bruce J Tonge, John L Bradshaw, Robert Iansek, Peter G Enticott, and Jenny McGinley. Gait function in high-functioning autism and asperger’s disorder. European child & adolescent psychiatry, 15(5):256–264, 2006.

[38] Juliane Rudeck, Silvia Vogl, Stefanie Banneke, Gilbert Schönfelder, and Lars Lewejohann. Repeatability analysis improves the reliability of behavioral data. PloS one, 15(4):e0230900, 2020.

[39] Paul R Sanberg, SA Zoloty, R Willis, CD Ticarich, K Rhoads, RP Nagy, SG Mitchell, AR Laforest, JA Jenks, LJ Harkabus, et al. Digiscan activity: automated measurement of thigmotactic and stereotypic behavior in rats. Pharmacology Biochemistry and Behavior, 27(3):569–572, 1987.

[40] Keith Sheppard, Justin Gardin, Gautam Sabnis, Asaf Peer, Megan Darrell, Sean Deats, Brian Geuther, Cathleen M. Lutz, and Vivek Kumar. Gait-level analysis of mouse open field behavior using deep learning-based pose estimation. bioRxiv, 2020.

[41] Jill L Silverman, Seda S Tolu, Charlotte L Barkan, and Jacqueline N Crawley. Repetitive self-grooming behavior in the btbr mouse model of autism is blocked by the mglur5 antagonist mpep. Neuropsychopharmacology, 35(4):976–989, 2010.

[42] Jill L Silverman, Sarah M Turner, Charlotte L Barkan, Seda S Tolu, Roheeni Saxena, Albert Y Hung, Morgan Sheng, and Jacqueline N Crawley. Sociability and motor functions in shank1 mutant mice. Brain research, 1380:120–137, 2011.

[43] Autism Speaks. Dsm-5 diagnostic criteria. New York: NY. Author retrieved, August, 10:2014, 2014.

[44] Kerri L Staples and Greg Reid. Fundamental movement skills and autism spectrum disorders. Journal of autism and developmental disorders, 40(2):209–217, 2010.

[45] Catherine J Stoodley, Anila M D’Mello, Jacob Ellegood, Vikram Jakkamsetti, Pei Liu, Mary Beth Nebel, Jennifer M Gibson, Elyza Kelly, Fantao Meng, Christopher A Cano, et al. Altered cerebellar connectivity in autism and cerebellar-mediated rescue of autism-related behaviors in mice. Nature neuroscience, 20(12):1744, 2017.

[46] Peter T Tsai, Court Hull, YunXiang Chu, Emily Greene-Colozzi, Abbey R Sadowski, Jarrett M Leech, Jason Steinberg, Jacqueline N Crawley, Wade G Regehr, and Mustafa Sahin. Autistic-like behaviour and cerebellar dysfunction in purkinje cell tsc1 mutant mice. Nature, 488(7413):647, 2012.

[47] Laurens Van der Maaten and Geoffrey Hinton. Visualizing data using t-sne. Journal of machine learning research, 9(11), 2008.

[48] Merina Varghese, Neha Keshav, Sarah Jacot-Descombes, Tahia Warda, Bridget Wicinski, Dara L Dickstein, Hala Harony-Nicolas, Silvia De Rubeis, Elodie Drapeau, Joseph D Buxbaum, et al. Autism spectrum disorder: neuropathology and animal models. Acta neuropathologica, 134(4):537–566, 2017.

[49] Roger N Walsh and Robert A Cummins. The open-field test: a critical review. Psychological bulletin, 83(3):482, 1976.

[50] Samuel S-H Wang, Alexander D Kloth, and Aleksandra Badura. The cerebellum, sensitive periods, and autism. Neuron, 83(3):518–532, 2014.

